# Rapid whole brain imaging of neural activity in freely behaving larval zebrafish (*Danio rerio*)

**DOI:** 10.1101/131532

**Authors:** Lin Cong, Zeguan Wang, Yuming Chai, Wei Hang, Chunfeng Shang, Wenbin Yang, Lu Bai, Jiulin Du, Kai Wang, Quan Wen

## Abstract

The internal brain dynamics that link sensation and action are arguably better studied during natural animal behaviors. Here we report on a novel volume imaging and 3D tracking technique that monitors whole brain neural activity in freely swimming larval zebrafish (*Danio rerio*). We demonstrated the capability of our system through functional imaging of neural activity during visually evoked and prey capture behaviors in larval zebrafish.

## Introduction

A central goal in systems neuroscience is to understand how distributed neural circuitry dynamics drive animal behaviors. The emerging field of optical neurophysiology allows the monitoring [1, 2] and manipulating [3-5] of the activities of defined populations of neurons that express genetically encoded activity indicators [6, 7] and light-activated proteins [1, 4, 5, 8]. Larval zebrafish (*Danio rerio*) are an attractive model system to investigate the neural correlates of behaviors owing to their small brain size, optical transparency, and rich behavioral repertoire [9, 10]. Whole brain imaging of larval zebrafish using light sheet/two-photon microscopy holds considerable potential in creating a comprehensive functional map that links neuronal activities and behaviors [11-13].

Recording neural activity maps in larval zebrafish has been successfully integrated with the virtual reality paradigm: closed-loop fictive behaviors in immobilized fish can be monitored and controlled via visual feedback that varies according to the electrical output patterns of motor neurons [11, 14]. The behavioral repertoire, however, may be further expanded in freely swimming zebrafish whose behavioral states can be directly inferred and when sensory feedback loops are mostly intact and active. For example, it is likely that vestibular as well as proprioceptive feedbacks are perturbed in immobilized zebrafish [14, 15]. The crowning moment during hunting behavior [16-18] — when a fish succeeds in catching a paramecium — cannot be easily replicated in a virtual reality setting. Therefore, whole brain imaging in a freely swimming zebrafish may allow optical interrogation of brain circuits underlying a range of less explored behaviors.

Although whole brain functional imaging methods are available for head-fixed larval zebrafish, imaging a speeding brain imposes many technical challenges. Current studies on freely swimming zebrafish are either limited to non-imaging optical systems [19] or wide field imaging at low resolution [20]. While light sheet microscopy (LSM) has demonstrated entire brain coverage and single neuron resolution in restrained zebrafish [12], it lacks the speed to follow rapid fish movement. Moreover, in LSM, the sample is illuminated from its side, a configuration that is difficult to be integrated with a tracking system. Conventional light field microscopy (LFM) [21, 22] is a promising alternative due to its higher imaging speed; however, its spatial resolution is relatively low. Specialized LFMs for monitoring neural activity utilizing temporal information were also developed recently [23, 24], which rely on spatiotemporal sparsity of fluorescent signals and cannot be applied to moving animals.

Here, we describe a fast 3D tracking technique and a novel volume imaging method that allow whole brain calcium imaging with high spatial and temporal resolution in freely behaving larval zebrafish. Zebrafish larvae possess extraordinary mobility. They can move at an instantaneous velocity up to 50 mm/s [25] and acceleration of 1 g (9.83 m/s^2^). To continuously track fish motion, we developed a high-speed closed-loop system in which (1) customized machine vision software allowed rapid estimation of fish movement in both the x-y and z directions; and, (2) feedback control signals drove a high-speed motorized x-y stage (at 300 Hz) and a piezo Z stage (at 100 Hz) to retain the entire fish head within the field of view of a high numerical aperture (25×, NA = 1.05) objective.

Larval zebrafish can make sudden and swift movements that easily cause motion blur and severely degrade imaging quality. To overcome this obstacle, we developed a new eXtended field of view LFM (XLFM). The XLFM can image sparse neural activity over the larval zebrafish brain at near single cell resolution and at a volume rate of 77 Hz, with the aid of genetically encoded calcium indicator GCamp6f. Furthermore, the implementation of flashed fluorescence excitation (200 μs in duration) allowed blur-free fluorescent images to be captured when a zebrafish moved at a speed up to 10 mm/s. The seamless integration of the tracking and imaging system made it possible to reveal rich whole brain neural dynamics during natural behavior with unprecedented resolution. We demonstrated the ability of our system during visually evoked and prey capture behaviors in larval zebrafish.

## Results

The newly developed XLFM is based on the general principle of light field [26] and can acquire 3D information from a single camera frame. XLFM greatly relaxed the constraint imposed by the tradeoff between spatial resolution and imaging volume coverage in conventional LFM. This achievement relies on optics and in computational reconstruction techniques. First, a customized lenslet array (Figure 1a, Figure 1-figure supplement 1) was placed at the rear pupil plane of the imaging objective, instead of at the imaging plane as in LFM. Therefore, in ideal conditions, a spatially invariant point spread function (PSF) could be defined and measured. In practice, the PSF was approximately spatially invariant, as discussed below. Second, the aperture size of each micro-lens was decoupled from their interspacing and spatial arrangement, so that both the imaging volume and the resolution could be optimized simultaneously given the limited imaging sensor size. Third, multifocal imaging [27, 28] was introduced to further increase the depth of view by dividing the micro-lenses array into several groups whose focal planes were at different axial positions (Figures 1b & c, Figure 1-figure supplements 3 & 4). Fourth, a new computational algorithm based on optical wave theory was developed to reconstruct the entire 3D volume from one image (Figure 1-figure supplement 5) captured by a fast camera (see Methods).

**Figure 1.**
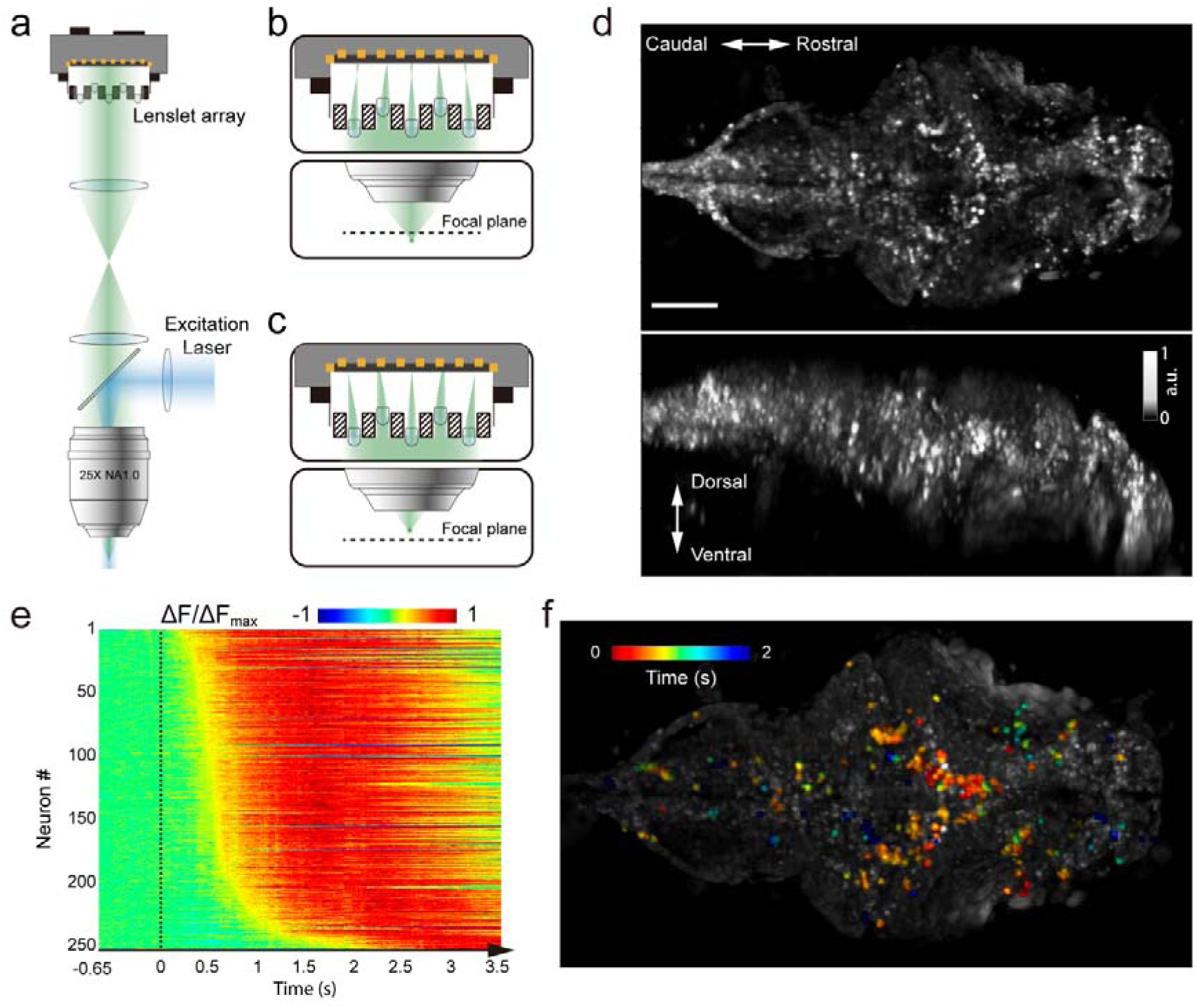
Whole brain imaging of larval zebrafish with XLFM. (a) Schematic of XLFM. Lenslet array position was conjugated to the rear pupil plane of the imaging objective. Excitation laser (blue) provided uniform illumination across the sample. (b–c) Point sources at two different depths formed, through two different groups of micro-lenses, sharp images on the imaging sensor, with positional information reconstructed from these distinct patterns. (d) Maximum intensity projections (MIPs) on time and space of time series volume images of an agarose-restrained larval zebrafish with pan-neuronal nucleus-localized GCaMP6f (huc:h2b-gcamp6f) fluorescence labeling. (e) Normalized neuronal activities of selected neurons exhibited increasing calcium responses after the onset of light stimulation at t = 0. Neurons were ordered by the onset time when the measured fluorescence signals reached 20% of their maximum. (f) Selected neurons in (e) were color coded based on their response onset time. Scale bar is 100 μm.

We characterized the XLFM by imaging 0.5 μm diameter fluorescent beads. In our design, the system had ~ Ø800 μm in plane coverage (Ø is the diameter of the lateral field of view) and more than 400 μm depth of view, within which an optimal resolution of 3.4 μm × 3.4 μm × 5 μm could be achieved over a depth of 200 μm (Figure 1-figure supplements 6 & 7, Methods) when sample was sparse. In the current implementation, however, the imaging performance suffered from variation in the focal length of the micro-lenses (Figure 1-figure supplement 8) and the optimal resolution at 3.4 μm × 3.4 μm × 5 μm was preserved over a reduced volume of Ø500 μm × 100 μm (Figure 1-figure supplements 9 & 10). Beyond this volume, the resolution degraded gradually. To minimize the reconstruction time while assuring whole brain coverage (~ 250 μm thick), all imaging reconstructions were carried out over a volume of Ø800 μm × 400 μm.

The achievable optimal resolution also relies on the sparseness of the sample, because the information captured by the image sensor was insufficient to assign independent values for all voxels in the entire reconstructed imaging volume. Given the total number of neurons (~ 80,000 [29]) in a larval zebrafish brain, we next introduced a sparseness index *ρ*, defined as the fraction of neurons in the brain active at a given instant, and used numerical simulation to characterize the dependence of achievable resolution on *ρ*. We identified a critical *ρ*_*c*_ ≈ 0.11, below which active neurons could be resolved at the optimal resolution (Figure 1-figure supplement 11b). As *ρ* increased, closely clustered neurons could no longer be well resolved (Figure 1-figure supplements 11c-d). Therefore, sparse neural activity is a prerequisite in XLFM for resolving individual neurons at the optimal resolution. Moreover, the above characterization assumed an aberration and scattering free environment; complex optical properties of biological tissue could also degrade the resolution [30].

We demonstrated the capabilities of XLFM by imaging the whole brain neuronal activities of a larval zebrafish (5 d post-fertilization (dpf)) at a speed of 77 volumes/s and relatively low excitation laser exposure of 2.5 mW/mm^2^ (Figure 1d, Video 1). The fluorescent intensity loss due to photobleaching reached ~ 50% when the zebrafish, which expressed pan-neuronal nucleus-labelled GCamp6f (huc:h2b-gcamp6f), was imaged continuously for ~ 100 min and over more than 300,000 volumes (Figure 1-figure supplement 12, Videos 2 & 3). To test whether XLFM could monitor fast changes in neuronal dynamics across the whole brain at high resolution (close to single neuron level), we first presented the larval zebrafish, restrained in low melting point agarose, with visual stimulation (~ 2.6 s duration). We found that different groups of neurons in the forebrain, midbrain, and hindbrain were activated at different times (Figures 1e–f, Videos 1 & 4), suggesting rapid sensorimotor transformation across different brain regions.

To track freely swimming larval zebrafish, we transferred fish into a water-filled chamber with a glass ceiling and floor. The 20 mm × 20 mm × 0.8 mm-sized chamber was coupled with a piezo actuator and mounted on a high-speed 2D motorized stage (Figure 2). A tracking camera monitored the lateral movement of the fish, and an autofocus camera, which captured light field images, monitored the axial movement of the fish head (Figure 2, Figure 2-figure supplement 1).

**Figure 2.**
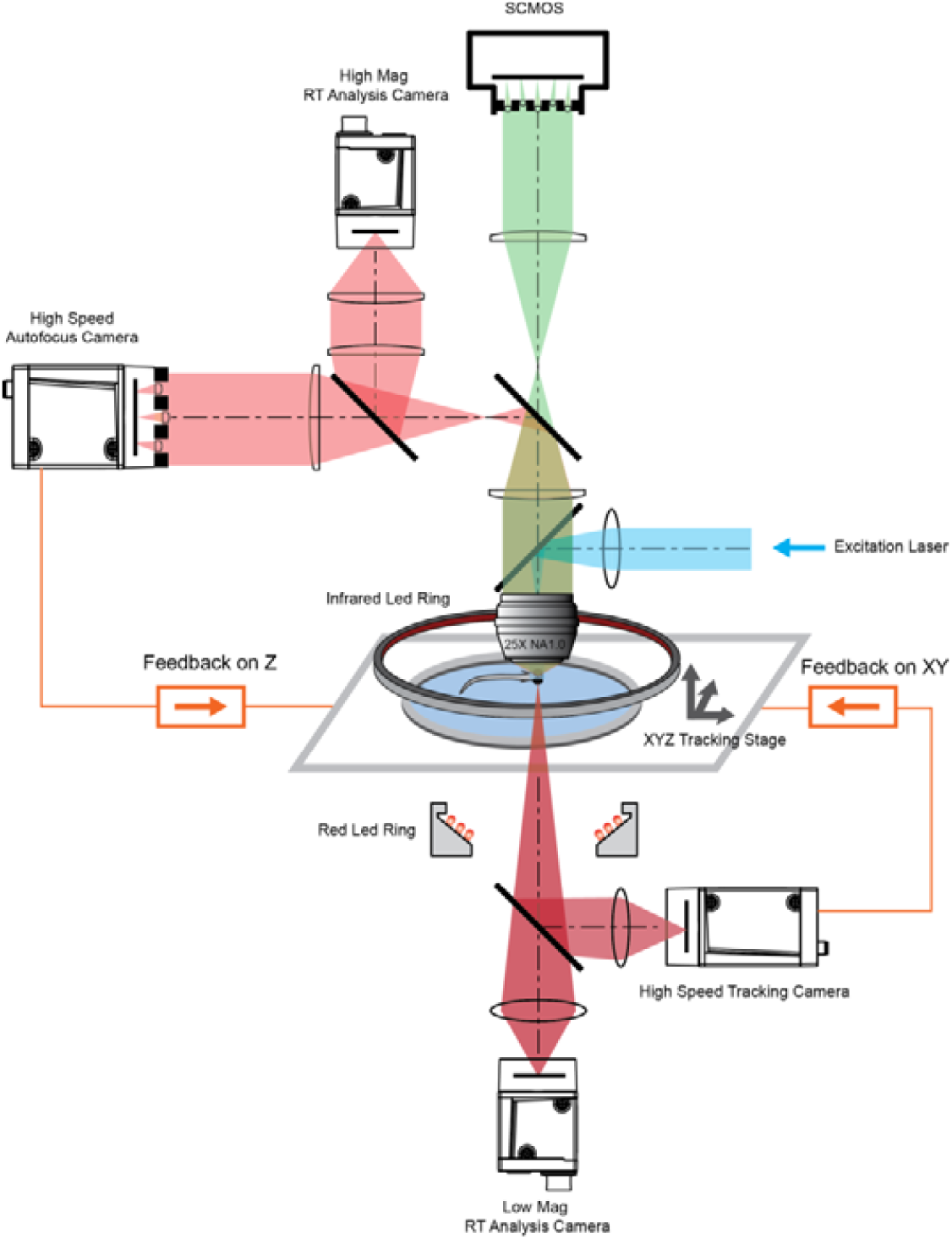
System schematics that integrated tracking, whole brain functional imaging, and real time behavioral analysis. Larval zebrafish swam in a customized chamber with an optically transparent ceiling and floor. The water-filled chamber was mounted on a high-speed three-axis stage (PI M686 & PI P725KHDS). Customized LED rings generated dark field illumination of the zebrafish. The scattered light was collected by four cameras: two cameras below the chamber were used for x-y plane tracking and low magnification real-time (RT) analysis, respectively; two cameras above the chamber and after the imaging objective were used for Z autofocus and high magnification RT analysis. The positional information of the larval zebrafish, acquired from the tracking and autofocus system, was converted to feedback voltage signals to drive the three-axis stage and to compensate for fish movement. The functional imaging system, described in Figure 1, shared the same imaging objective placed above the swimming chamber. The 3D tracking, RT behavioral analysis, and functional imaging system were synchronized for accurate correlation between neural activity and behavioral output.

Real-time machine vision algorithms allowed quick estimate of lateral (within 1 ms) and axial (~ 5 ms) head positions (see Methods). The error signals in three dimensions, defined as the difference between the head position and set point, were calculated (Figure 3a) and converted to analog voltage signals through proportional-integral-derivative (PID) control to drive the motorized stage and z-piezo scanner. Tracking and autofocusing allowed for rapid compensation of 3D fish movement (300 Hz in x and y, 100 Hz in z, Figure 3a) and retainment of the fish head within the field of view of the imaging objective.

**Figure 3.**
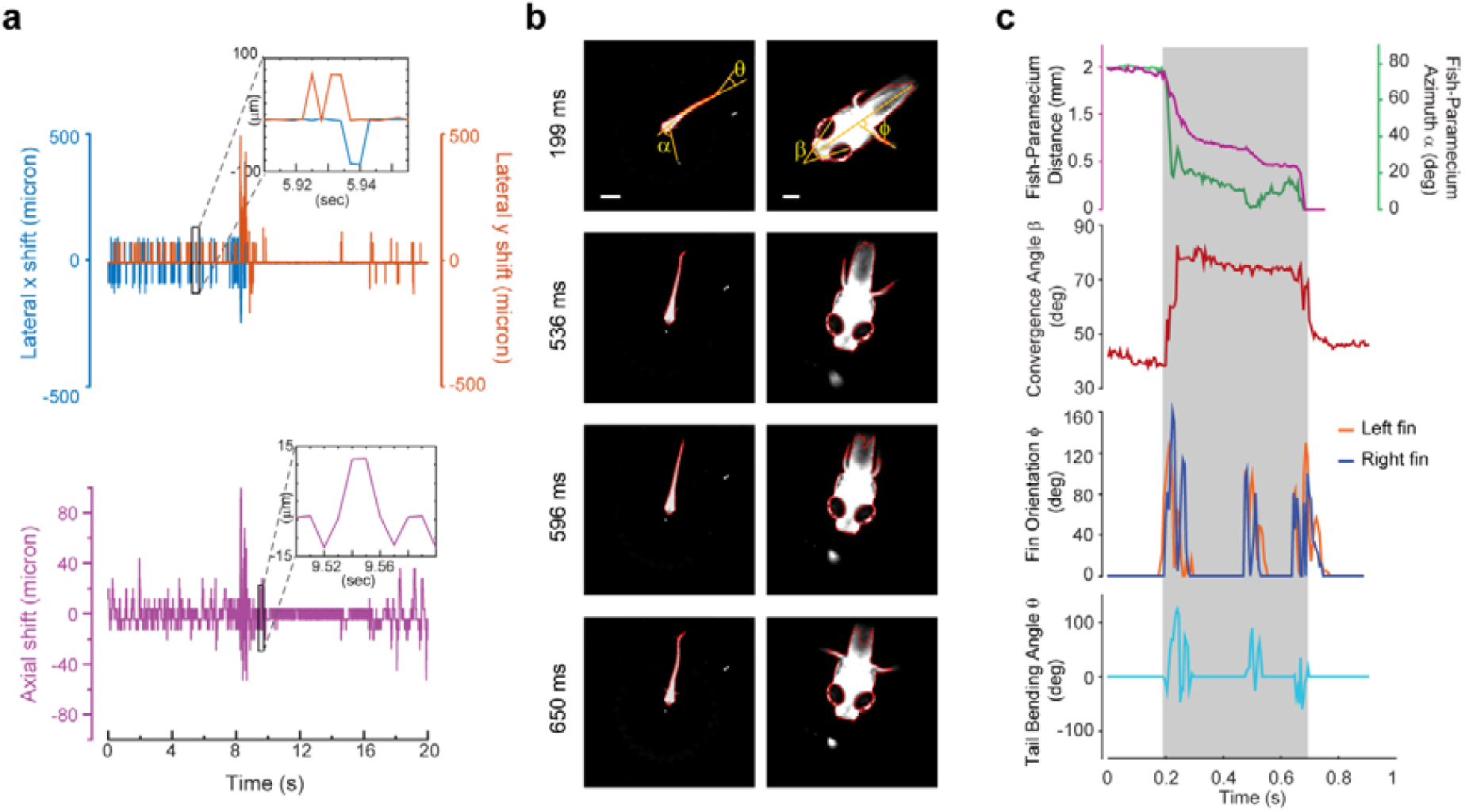
3D tracking of larval zebrafish. (a) Representative time varying error signals in three dimensions, defined as the difference between real head position and set point. Inset provides magnified view at short time interval. Lateral movement can be rapidly compensated for within a few milliseconds with an instantaneous velocity of up to 10 mm/s. The axial shift was small compared with the depth coverage (200 μm) during whole brain imaging, and thereby had minor effect on brain activity reconstruction. (b) Tracking images at four time points during prey capture behavior, acquired at low (left) and high (right) magnification simultaneously. Scale bars are 1 mm (left) and 200 μm (right). (c) Kinematics of behavioral features during prey capture. Shaded region marks the beginning and end of the prey capture process.

Our tracking system permitted high-speed and high-resolution recording of larval zebrafish behaviors. With two cameras acquiring head and whole body videos simultaneously (Figure 2, Figure 3b), we recorded and analyzed in real time (see Methods) the kinematics of key features during larval zebrafish prey capture (Figures 3b & c, Videos 5 & 6). Consistent with several earlier findings [16-18], eyes converged rapidly when the fish entered the prey capture state (Figure 3c). Other features that characterized tail and fin movement were also analyzed at high temporal resolution (Figure 3c).

The integration of the XLFM and 3D tracking system allowed us to perform whole brain functional imaging of a freely behaving larval zebrafish (Figure 2). We first replicated the light-evoked experiment (similar to Figure 1), albeit in a freely behaving zebrafish with pan-neuronal cytoplasm-labeled GCaMP6s (huc:gcamp6s), which exhibited faster and more prominent calcium response (Video 7). Strong activities were observed in the neuropil of the optical tectum and the midbrain after stimulus onset. The fish tried to avoid strong light exposure and made quick tail movement at ~ 60 Hz. Whole brain neural activity was monitored continuously during the light-evoked behavior, except for occasional blurred frames due to the limited speed and acceleration of the tracking stage.

Next, we captured whole brain neural activity during the entire prey capture process in freely swimming larval zebrafish (huc:gcamp6s, Video 8). When a paramecium moved into the visual field of the fish, groups of neurons, indicated as group 1 in Figure 4b, near the contralateral optical tectum of the fish were first activated (t_1_). The fish then converged its eyes onto the paramecium and changed its heading direction in approach (t_2_). Starting from t_2_, several groups of neurons in the hypothalamus, midbrain, and hindbrain, highlighted as groups 2, 3, and 4 in Figure 4b, were activated. It took the fish three attempts (Figure 4c) to catch and eat the paramecium. After the last try (t_4_), neuron activity in group 1 decreased gradually, whereas activities in the other groups of neurons continued to rise and persisted for ~ 1 s before the calcium signals decreased. The earliest tectal activity (group 1) responsible for prey detection found here is consistent with previous studies [31, 32]. Moreover, our data revealed interesting neural dynamics arising from other brain regions during and after successful prey capture. We also monitored similar behavior in a zebrafish expressing nucleus-localized GCamp6f (huc:h2b-gcamp6f) with better resolution but less prominent calcium response (Video 9).

**Figure 4.**
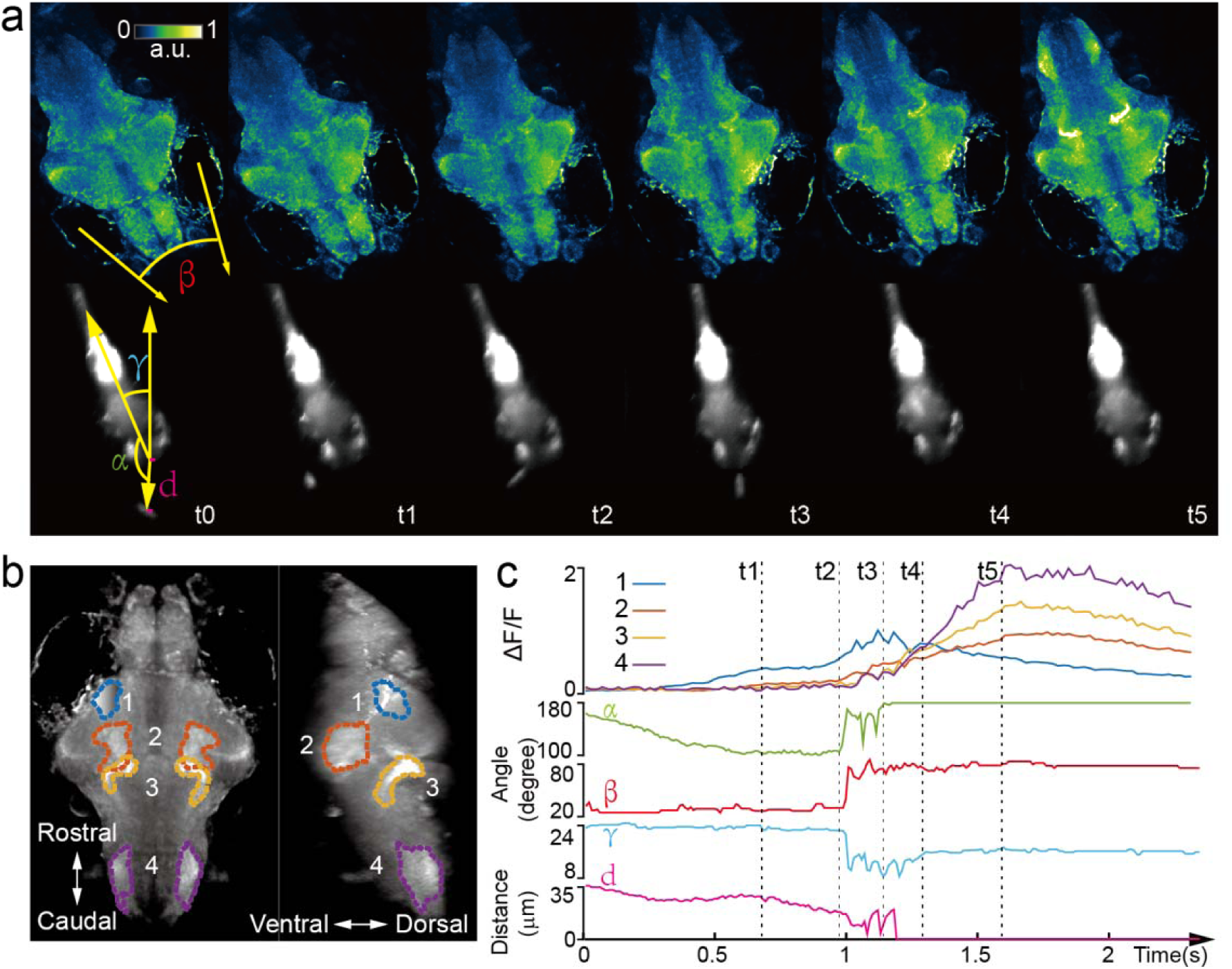
Whole brain imaging of larval zebrafish during prey capture behavior. (a) Renderings of whole brain calcium activity at six time points (up) and the corresponding behavioral images (bottom). Features used to quantify behavior were: fish-paramecium azimuth *α*; convergence angle between eyes β; head orientation γ; and fish-paramecium distance *d*. (b) Maximum intensity projections of zebrafish brain with pan-neuronal cytoplasm-labeled GCaMP6s (huc:gcamp6s). Boundaries of four brain regions are color marked. (c) Neural dynamics inferred from GCaMP6 fluorescence changes in these four regions during the entire prey capture behavior (up) and the kinematics of behavioral features (bottom). Note that between t2 and t4, fish-paramecium distance *d* exhibits three abrupt kinks, representing the three attempts to catch prey.

## Discussion

Whole brain imaging in freely behaving animals has been previously reported in *Caenorhabditis elegans*, by integrating spinning-disk confocal microscopy with a 2D tracking system [33, 34]. In the more remote past, Howard Berg pioneered the use of 3D tracking microscopy to study bacteria chemotaxis [35]. However, the significant increase of animal size imposes challenges both in tracking and imaging technologies. The XLFM, derived from the general concept of light field imaging [21, 26, 36, 37], overcomes several critical limitations of conventional LFM and allows optimization of imaging volume, resolution, and speed simultaneously. Furthermore, it can be perfectly combined with flashed fluorescence excitation to capture blur-free images at high resolution during rapid fish movement. Taken together, we have developed a volume imaging and tracking microscopy system suitable for observing and capturing freely behaving larval zebrafish, which have ~ 80,000 neurons and can move two orders of magnitude faster than *C. elegans*.

Tracking and whole brain imaging of naturally behaving zebrafish provide an additional way to study sensorimotor transformation across the brain circuit. A large body of research suggests that sensory information processing depends strongly on the locomotor state of an animal [38-40]. The ability to sense self-motion, such as proprioceptive feedback [41] and efferent copy [42], can also profoundly shape the dynamics of the neural circuit and perception. To explore brain activity in swimming zebrafish, several studies have utilized an elegant tail-free embedding preparation [25, 43, 44], in which only the head of the fish is restrained in agarose for functional imaging. Nevertheless, it would be ideal to have physiological access to all neurons in defined behavioral states, where all sensory feedback loops remain intact and functional. Our XLFM-3D tracking system is one step towards this goal, and could be better exploited to explore the neural basis of more sophisticated natural behaviors, such as prey capture and social interaction, where the integration of multiple sensory feedbacks becomes critical.

In the XLFM, the camera sensor size limited the number of voxels and hence the number of neurons that could be reliably reconstructed. Our simulation suggested that the sparseness of neuronal activities is critical for optimal imaging volume reconstruction. A growing body of experimental data indeed suggests that population neuronal activities are sparse [45, 46] and sparse representation is useful for efficient neural computation [47, 48]. Given the total number of neurons in the larval zebrafish brain, we found that when the fraction of active neurons in a given imaging frame was less than *ρ*_*c*_ ≈ 0.11, individual neurons could be resolved at optimal resolution. When population neural activity was dense (*e*.*g*., neurons have high firing rate and firing patterns have large spatiotemporal correlation), we obtained a coarse-grained neural activity map with reduced resolution.

To retain the fish head within the field of view of the imaging objective, our tracking system compensated for fish movement by continuously adjusting the lateral positions of the motorized stage. As a result, self-motion perceived by the fish was not exactly the same as that during natural behaviors. The linear acceleration of the swimming fish, encoded by vestibular feedback, was significantly underestimated. The perception of angular acceleration during head orientation remained largely intact. The relative flow velocity along the fish body, which was invariant upon stage translation, can still be detected by specific hair cells in the lateral line system [49, 50]. Together, the interpretation of brain activity associated with self-motion must consider motion compensation driven by the tracking system.

Both tracking and imaging techniques can be improved in the future. For example, the current axial displacement employed by the piezo scanner had a limited travelling range (400 μm), and our swimming chamber essentially restrained the movement of the zebrafish in two dimensions. This limitation could be relaxed by employing axial translation with larger travelling range and faster dynamics. Furthermore, to avoid any potential disturbance of animal behaviors, it would be ideal if the imaging system moved, instead of the swimming chamber.

In XLFM, the performance degradation caused by focal length variation of the micro-lenses could be resolved by higher precision machining. In addition, the capability of XLFM could be further improved with the aid of technology development in other areas. With more pixels in the imaging sensor, we could resolve more densely labelled samples, and achieve higher spatial resolution without sacrificing imaging volume coverage by introducing more than two different focal planes formed by more groups of micro-lenses. With better imaging objectives that could provide higher numerical aperture and larger field of view at the same time, we could potentially image the entire nervous system of the larval zebrafish with single neuron resolution in all three dimensions. Additionally, the fast imaging speed of XLFM holds the potential for recording electrical activity when high signal-to-noise ratio (SNR) fluorescent voltage sensors become available [51]. Finally, the illumination-independent characteristic of XLFM is perfectly suitable for recording brain activities from bioluminescent calcium/voltage indicators in a truly natural environment, where light interference arising from fluorescence excitation can be eliminated [19].

## METHODS

### XLFM

The imaging system (Figure 1) was a customized upright microscope. Along the fluorescence excitation light path, a blue laser (Coherent, OBIS 488 nm, 100 mW, USA) was expanded and collimated into a beam with a diameter of ~ 25 mm. It was then focused by an achromatic lens (focal length: 125 mm) and reflected by a dichroic mirror (Semrock, Di02-R488-25x36, USA) into the back pupil of the imaging objective (Olympus, XLPLN25XWMP2, 25X, NA 1.05, WD 2mm, Japan) to result in an illumination area of ~1.44 mm in diameter near the objective’s focal plane. In the fluorescence imaging light path, excited fluorescence was collected by the imaging objective and transmitted through the dichroic mirror. A pair of achromatic lenses (focal lengths: F1 = 180 mm & F2 = 160 mm), arranged in 2F1 + 2F2, were placed after the objective and dichroic mirror to conjugate the objective’s back pupil onto a customized lenslet array (Figure 1-figure supplement 1). The customized lenslet array was an aluminum plate with 27 holes (1.3 mm diameter aperture on one side and 1 mm diameter aperture on the other side, Source Code File 1) housing 27 customized micro-lenses (1.3 mm diameter, focal length: 26 mm). The 27 micro-lenses were divided into two groups (Figure 1-figure supplement 1) and an axial displacement of 2.5 mm was introduced between them. Due to the blockage of light by the aluminum micro-lenses housing, 16% of the light after a 1.05 NA imaging objective was effectively collected by the camera. This efficiency is equivalent to using a 0.4 NA imaging objective. Finally, the imaging sensor of a sCMOS camera (Hamamatsu, Orca-Flash 4.0 v2, Japan) was placed at the middle plane between two focal planes formed by two different groups of micro-lenses. The total magnification of the imaging system was ~ 4, so one camera pixel (6.5 μm) corresponded to ~1.6 μm on the sample.

We developed a computational algorithm for 3D volume reconstruction, which required an accurately measured PSF (Figure 1-figure supplement 2). The PSF was measured by recording images of a 500 nm diameter fluorescent bead sitting on a motorized stage under the objective. A stack of 200 images was recorded when the bead was scanned with a step size of 2 μm in the axial direction from 200 μm below the objective’s focal plane to 200 μm above. Since the images formed by two different groups of micro-lenses were from different axial locations and had different magnifications, the measured raw PSF data were reorganized into two complementary parts: PSF_A and PSF_B (Figure 1-figure supplements 3 & 4), according to the spatial arrangement of the micro-lenses. We took PSF_A stack, PSF_B stack, and a single frame of a raw image (2048 × 2048 pixels) as inputs, and applied a newly developed algorithm to reconstruct the 3D volume.

### Image reconstruction of XLFM

The reconstruction algorithm was derived from the Richardson-Lucy deconvolution. The goal was to reconstruct a 3D fluorescent object from a 2D image:

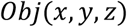

The algorithm assumes that the real 3D object can be approximated by a discrete number of x-y planes at different *z* positions:

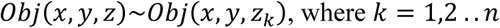

The numbers and positions of these planes can be arbitrary, yet the Nyquist sampling rate should be chosen to optimize the speed and accuracy of the reconstruction.

As the imaging system consisted of two different groups of micro-lenses (Figure 1-figure supplement 1), their PSFs (Figure 1-figure supplements 3 & 4) each consisted of a stack of planes that were measured at the same chosen axial positions *z*_*k*_:

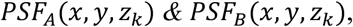

Although the PSF was measured in imaging space, here we denote *x* and *y* as coordinates in object space to follow conventions in optical microscopy. Here and below, the combination of *PSF*_*A*_ and *PSF*_*B*_ is the total PSF.

Additionally, the images formed by two different groups of micro-lenses had different magnifications, which could be determined experimentally. The ratio between two different magnifications can be defined as:

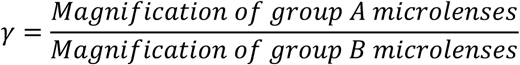

Then the captured image on the camera can be estimated as:

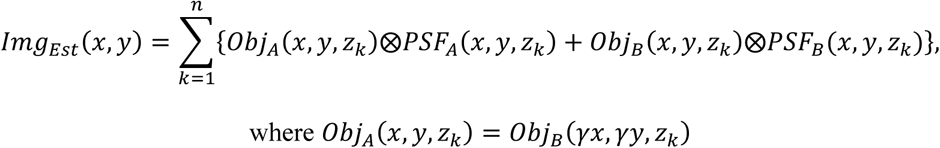

The operator ⨂ represents 2D convolution. Here, *x* and *y* on the left hand side of the equation also represent coordinates in object space so that 2D convolution was carried out in the same coordinates.

The goal of the algorithm is to estimate the *Obj*(*x*, *y*, *z*_*k*_) from the measured camera frame:

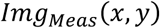

According to the Richardson-Lucy deconvolution algorithm, the iterative reconstruction can be expressed as:

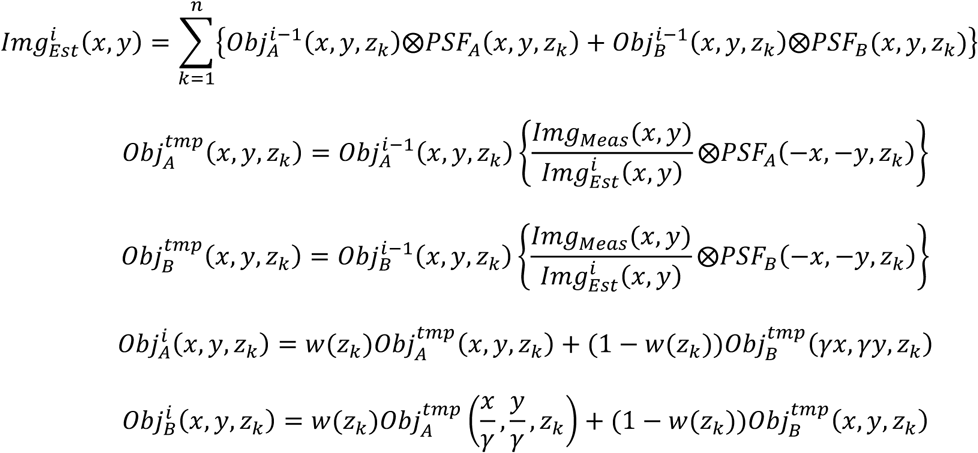

Here 0 ≤ *w*(*z*_*k*_) ≤ 1 is the weighting factor at different axial positions. The choice of *w*(*z_k_*) can be arbitrary. Because the resolutions achieved by different groups of micro-lenses at different z positions were not the same, the weighting factor can take this effect into consideration by weighing higher quality information more than lower quality information. One simple choice is *w*(*z*_*k*_) = 0.5, that is, to weigh information from two groups of micro-lenses equally.

The starting estimate of the object can be any non-zero value. Near the end of the iterations, 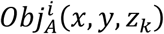 and 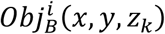 are interchangeable, except with different magnifications. Either can be used as the resulting estimate of the 3D object.

In XLFM, together with its reconstruction algorithm, the diffraction of the 3D light field is properly considered by experimentally measured PSF. The raw imaging data can be fed into the algorithm directly without any preprocessing. Given that the PSF is spatially invariant, which is satisfied apart from small aberrations, the algorithm can handle overlapping fish images (Figure 1-figure supplement 5). As a result, the field of view can be increased significantly. The reconstruction algorithm was typically terminated after 30 iterations when modifications in the estimated object became very small. The computation can speed up significantly via GPU. It took about 4 min to reconstruct one 3D volume using a desktop computer with a GPU (Nvidia Titan X). In comparison, the reconstruction ran ~20× slower using a CPU (Intel E5-2630v2) on a Dell desktop. The source code written in MATLAB can be found in the Source Code File 2.

The 3D deconvolution method has been developed for conventional LFM [21]. Our method differs from [21] in several ways. (1) The optical imaging systems are different. (2) The definitions of PSFs are different. Ours defines a spatially *invariant* PSF (see below for detailed characterization), whereas [21] defined a spatially variant PSF, leading to increased computational complexity in the deconvolution algorithm. (3) The PSF in [21] was simulated based on a model derived from an ideal imaging system, whereas ours was measured experimentally. Furthermore, our system took practical conditions, such as a non-ideal imaging objective, actual positions of microlenses, the spectrum of received fluorescence signal *et al*., into consideration.

### Resolution characterization of XLFM

Unlike conventional microscopy, where the performance of the imaging system is fully characterized by the PSF at the focal plane, the capability of XLFM is better characterized as a function of positions throughout the imaging volume.

We first characterized the spatial resolution in the x-y plane by analyzing the spatial frequency support of the experimentally measured PSF from individual micro-lenses using a 0.5 μm diameter fluorescent bead. The optical transfer function (OTF), which is the Fourier transform of the PSF in the x-y plane, was extended to a spatial frequency of ~1/3.4 μm^−1^ (Figure 1-figure supplement 6), a result that agreed well with the designed resolution at 3.4 μm, given that the equivalent NA of individual micro-lenses was 0.075.

The lateral resolution, measured from the raw PSF behind individual micro-lenses, was preserved across the designed cylindrical imaging volume of Ø800 μm × 200 μm (Figure 1-figure supplement 6). However, the reconstruction results (Figure 1-figure supplement 9), which used total PSF (Figure 1-figure supplement 2), exhibited resolution degradation when the fluorescent bead was placed more than 250 μm away from the center (Figure 1-figure supplement 9). This discrepancy resulted from the variation in focal length of the micro-lenses (Figure 1-figure supplement 8), which, in turn, led to spatial variance of the defined *PSF*_*A*_ and *PSF*_*B*_. In principle, the designed lateral resolution of 3.4 μm could be preserved over a volume of Ø800 μm × 200 μm by reducing focal length variation to below 0.3%

We next characterized the axial resolution of the XLFM. The XLFM gained axial resolution by viewing the object from large projection angles achieved by micro-lenses sitting near the edge of the objective’s back pupil plane. For example, if two points of light source were located at the same position in the X-Y plane, but were separated by Δ*z* in the axial direction, then one micro-lens in the XLFM could capture an image of these two points with a shift between them. The shift can be determined as:

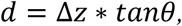

where *θ* is the inclination angle inferred from the measured PSF (Figure 1-figure supplement 2). If the two points in the image can be resolved, the two points separated by Δ*z* can be resolved by the imaging system. Since a micro-lens sitting in the outer layer of the array offered the largest inclination angle of 40 degree in our system, the axial resolution dz can be directly calculated as:

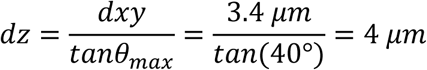

The best way to confirm the theoretical estimate is to image two fluorescent beads with precisely controlled axial separations. However, this is technically very challenging. Instead, we pursued an alternative method that is equivalent to imaging two beads simultaneously:

1. We took a z stack of images of fluorescent beads, as done in measuring the PSF.
2. In post processing, we added two images from different z positions to mimic the beads being present simultaneously at two different *z* positions.

The above method allowed us to experimentally characterize the axial resolution afforded by individual micro-lenses focusing at different z positions. We used a single fluorescent bead (0.5 μm in diameter) with a high SNR (Figure 1-figure supplement 7a). We imaged at different axial positions: *z* = −100 μm, *z* = 0 μm, and *z* = 100 μm (Figure 1-figure supplement 7b). The third column is the combined images in column 1 & 2. The capability of resolving the two beads in the third column can be demonstrated by spatial frequency analysis (fourth column in Figure 1-figure supplement 7b). The two line dips, indicating the existence of two beads instead of one rod in the fourth column, were confirmations of the resolving capability. This becomes more evident after deconvolution of the raw images (fifth column in Figure 1-figure supplement 7b). Micro-lenses 1 and 2 could resolve two beads, separated by 5μm, within the range of −100 μm ≤ *z* ≤ 0 and 0 ≤ *z* ≤ 100 μm, respectively. In other words, the complementary information provided by the two micro-lenses allowed the system to maintain a high axial resolution at 5 μm across a 200 μm depth.

Next, we imaged densely packed fluorescent beads (0.5 μm in diameter) with a low SNR (Figure 1-figure supplement 10a), and used our reconstruction algorithm to determine the minimum axial separation between beads that could be resolved (Figure 1-figure supplements 10b–c). In this case, 5 μm axial resolution could be preserved across a depth of 100 μm. The resolution decayed gradually to ~10 μm at the edge of an imaging volume with a 400 μm axial coverage (Figure 1-figure supplement 10b). We believe that the optimal axial resolution at 5 μm could be achieved over an axial coverage of 200 μm by minimizing micro-lens focal length variation (Figure 1-figure supplement 8).

Finally, we characterized how the imaging performance depended upon the sparseness of the sample. Given the total number of neurons (~ 80,000) in a larval zebrafish brain, we introduced a sparseness index *ρ*, defined as the fraction of neurons in the brain active at an imaging frame, and used numerical simulation to characterize the dependence of achievable resolution on *ρ*. To this end, we simulated a zebrafish larva with uniformly distributed firing neurons (red dots in Figure 1-figure supplement 11a). By convolving the simulated zebrafish with the experimentally measured PSFs (Figure 1-figure supplements 3 & 4), we generated an image that mimicked the raw data captured by the camera. We then reconstructed the simulated neurons from this image, represented by green dots. When *ρ* was equal to or less than 0.11, which corresponded to ~ 9000 neurons activated at a given instant, all active neurons, including those closely clustered, could be reconstructed with optimal resolution (Figure 1-figure supplement 11b inset). As the sparseness index *ρ* increased, the resolution degraded: nearby neurons merged laterally and elongated axially (Figure 1-figure supplements 11c–d). In all calculations, the Poisson noise was properly considered by assuming that each active neuron emitted 20,000 photons, 2.2% of which were collected by our imaging system.

*In vivo* resolution characterization is challenging due to a lack of bright and spot-like features in living animals. Additionally, achievable resolution depends on the optical properties of biological tissues, which can be highly heterogeneous and difficult to infer. The light scattering and aberration induced by biological tissue usually leads to degraded imaging performance [30, 52-54].

### XY tracking system

To compensate for lateral fish movement and retain the entire fish head within the field of view of a high NA objective (25×, NA = 1.05), a high-speed camera was used to capture fish motion (2 ms exposure time, 300 fps or higher, Basler aca2000-340kmNIR, Germany). We developed an FPGA-based RT system in LabVIEW that could rapidly identify the head position by processing the pixel stream data within the Cameralink card before the whole image was transferred to RAM. The error signal between the actual head position and the set point was then fed into the PID to generate output signals and control the movement of a high-speed motorized stage (PI M687 ultrasonic linear motor stage, Germany). In the case of large background noise, we alternatively performed conventional imaging processing in C/C++ (within 1 ms delay). The rate-limiting factor of our lateral tracking system was the response time of the stage (~ 300 Hz).

### Autofocus system

We applied the principle of LFM to determine the axial movement of larval zebrafish. The autofocus camera (100 fps, Basler aca2000-340kmNIR, Germany) behind a one-dimensional micro-lens array captured triplet images of the fish from different perspectives (Figure 2-figure supplement 1a). Z motion caused an extension or contraction between the centroids of the fish head in the left and right sub-images, an inter-fish distance (Figure 2-figure supplement 1b) that can be accurately computed from image autocorrelation. The inter-fish distance, multiplied by a pre-factor, can be used to estimate the z position of the fish, as it varies linearly with axial movement (Figure 2-figure supplement 1c). The error signal between the actual axial position of the fish head and the set point was then fed into the PID to generate an output signal to drive a piezo-coupled fish container. The feedback control system was written in LabVIEW. The code was further accelerated by parallel processing and the closed loop delay was ~ 5 ms. The rate-limiting factor of the autofocus system was the settling time of the piezo scanner (PI P725KHDS, Germany, 400 μm travelling distance), which was about 10 ms.

### Real-time behavioral analysis

Two high-speed cameras acquired dark-field images at high and low magnification, respectively, and customized machine vision software written in C/C++ with the aid of OpenCV library was used to perform real-time behavioral analysis of freely swimming larval zebrafish. At high magnification, eye positions, their orientation, and convergence angle were computed; at low magnification, the contour of the whole fish, centerline, body curvature, and bending angle of the tail were computed. The high mag RT analysis was run at ~ 120 fps and the low mag RT analysis was run at ~ 180 fps. The source code can be found in the Source Code File 3.

### Ethics statement and animal handling

All animal handling and care were conducted in strict accordance with the guidelines and regulations set forth by the Institute of Neuroscience, Chinese Academy of Sciences, University of Science and Technology of China (USTC) Animal Resources Center, and University Animal Care and Use Committee. The protocol was approved by the Committee on the Ethics of Animal Experiments of the USTC (permit number: USTCACUC1103013).

All larval zebrafish (huc:h2b-gcamp6f and huc:gcamp6s) were raised in embryo medium under 28.5°C and a 14/10 h light/dark cycle. Zebrafish were fed with paramecium from 4 dpf. For restrained experiments, 4–6 dpf zebrafish were embedded in 1% low melting point agarose. For freely moving experiments, 7–11 dpf zebrafish with 10% Hank’s solution were transferred to a customized chamber (20 mm in diameter, 0.8 mm in depth), and 10–20 paramecia were added before the chamber was covered by a coverslip.

### Neural activity analysis

To extract neural activity induced by visual stimuli (Figures 1e & f), time series 3D volume stacks were first converted to a single 3D volume stack, in which each voxel represented variance of voxel values over time. Candidate neurons were next extracted by identifying local maxima in the converted 3D volume stack. The region-of-interest (ROI) was set according to the empirical size of a neuron. The voxels around the local maxima were selected to represent neurons. The fluorescence intensity over each neuron’s ROI was integrated and extracted as neural activity. Relative fluorescent changes Δ*F*/*F*_0_ were normalized to their maximum calcium response Δ*F_max_*/*F*_0_ over time, and sorted according to their onset time when Δ*F* first reached 20% of its Δ*F*_*max*_ (Figures 1e & f) after the visual stimulus was presented.

### Visual stimulation

A short wavelength LED was optically filtered (short-pass optical filter with cut-off wavelength at 450 nm, Edmund #84-704) to avoid light interference with fluorescence. It was then focused by a lens into a spot 2~3 mm in diameter. The zebrafish was illuminated from its side. The total power of the beam was roughly 3 mW.

### Statement of replicates and repeats in experiments

Each experiment was repeated at least three times with similar experimental conditions. Imaging and video data acquired from behaviorally active larval zebrafish with normal huc:h2b-gcamp6f or huc:gcamp6s expression were used in the main figures and videos.

## Acknowledgements

We thank Misha B. Ahrens for the zebrafish lines. We thank Yong Jiang, Tongzhou Zhao, WenKai Han, Shenqi Fan for assistance in building the 3D tracking system, real time behavioral analysis, and larval zebrafish experiments. We thank Dr. Bing Hu and Dr. Jie He for his support in zebrafish handling and helpful discussions.

## Videos

**Video 1| Whole brain functional imaging of larval zebrafish under light stimulation** Whole brain XLFM imaging of a 5 dpf agarose-embedded larval zebrafish expressing nucleus-localized GCamp6f (huc:h2b-gcamp6f). Light stimulation was introduced at time point t = 0. Whole brain activity was recorded at 77 volumes/s.

**Video 2| Whole brain functional imaging of spontaneous activities of larval zebrafish** Whole brain XLFM imaging of a 5 dpf agarose-embedded larval zebrafish expressing nucleus-localized GCamp6f (huc:h2b-gcamp6f). Spontaneous neural activity was recorded at 0.6 volumes/s.

**Video 3| Whole brain functional imaging of spontaneous activities of larval zebrafish** Whole brain XLFM imaging of a 5 dpf agarose-embedded larval zebrafish expressing cytoplasm-labeled GCamp6s (huc:gcamp6s). Spontaneous neural activity was recorded at 0.6 volumes/s.

**Video 4| Whole brain functional imaging of larval zebrafish under light stimulation** Whole brain XLFM imaging of a 5 dpf agarose-embedded larval zebrafish expressing cytoplasm-labeled GCamp6s (huc:gcamp6s). Light stimulation was introduced at time point t = 0. Whole brain activity was recorded at 50 volumes/s.

**Video 5| Tracking of larval zebrafish during prey capture behavior at low resolution** Tracking and real time kinematic analysis of larval zebrafish during prey capture behavior at low resolution. Recorded at 190 frames/s.

**Video 6| Tracking of larval zebrafish during prey capture behavior at high resolution** Tracking and real time kinematic analysis of larval zebrafish during prey capture behavior at high resolution. Recorded at 160 frames/s.

**Video 7| Whole brain functional imaging of a freely swimming larval zebrafish under light stimulation** Whole brain XLFM imaging of a 7 dpf freely swimming larval zebrafish expressing cytoplasm-labeled GCamp6s (huc:gcamp6s). Light stimulation was introduced at time point t = 0. Whole brain activities were recorded at 77 volumes/s and with a flashed excitation laser under 0.3 ms exposure time.

**Video 8| Whole brain functional imaging of a freely swimming larval zebrafish during prey capture behavior** Whole brain XLFM imaging of an 11 dpf freely swimming larval zebrafish expressing cytoplasm-labeled GCamp6s (huc:gcamp6s). The entire process during which the larval zebrafish caught and ate the paramecium was recorded.

**Video 9| Whole brain functional imaging of a freely swimming larval zebrafish during prey capture behavior** Whole brain XLFM imaging of a 7 dpf freely swimming larval zebrafish expressing nucleus-localized GCamp6f (huc:h2b-gcamp6f). The entire process during which the larval zebrafish caught and ate the paramecium was recorded.

**Figure 1-figure supplement 1.**
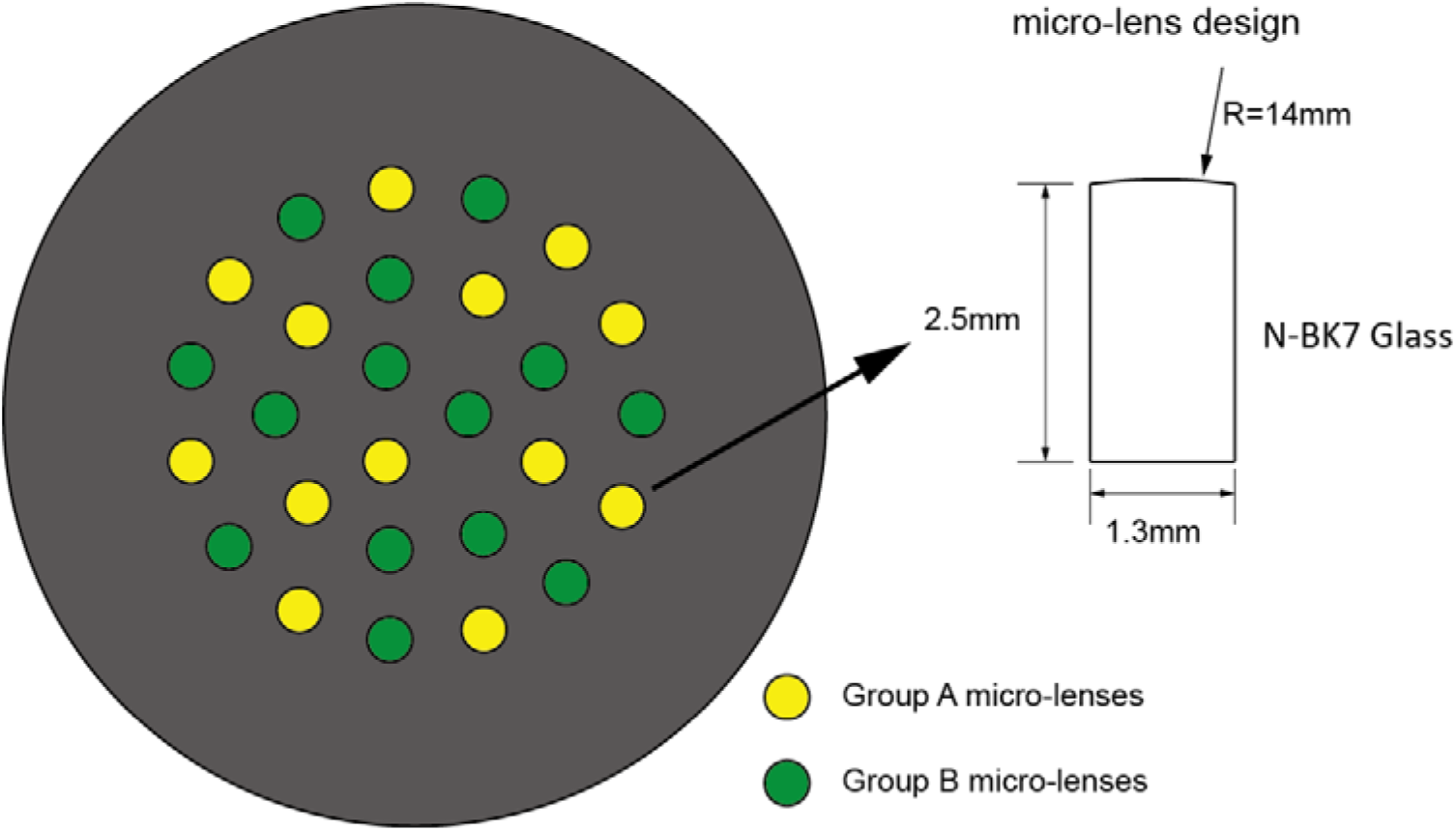
Customized lenslet array. Customized lenslet array consisted of 27 customized micro-lenses (1.3 mm diameter, 26 mm focal length) embedded in an aluminum plate with 27 drilled holes (1.3 mm diameter aperture on one side and 1 mm diameter aperture on the other side). Micro-lenses were divided into two groups (A or B), illustrated in yellow and green, respectively.

**Figure 1-figure supplement 2.**
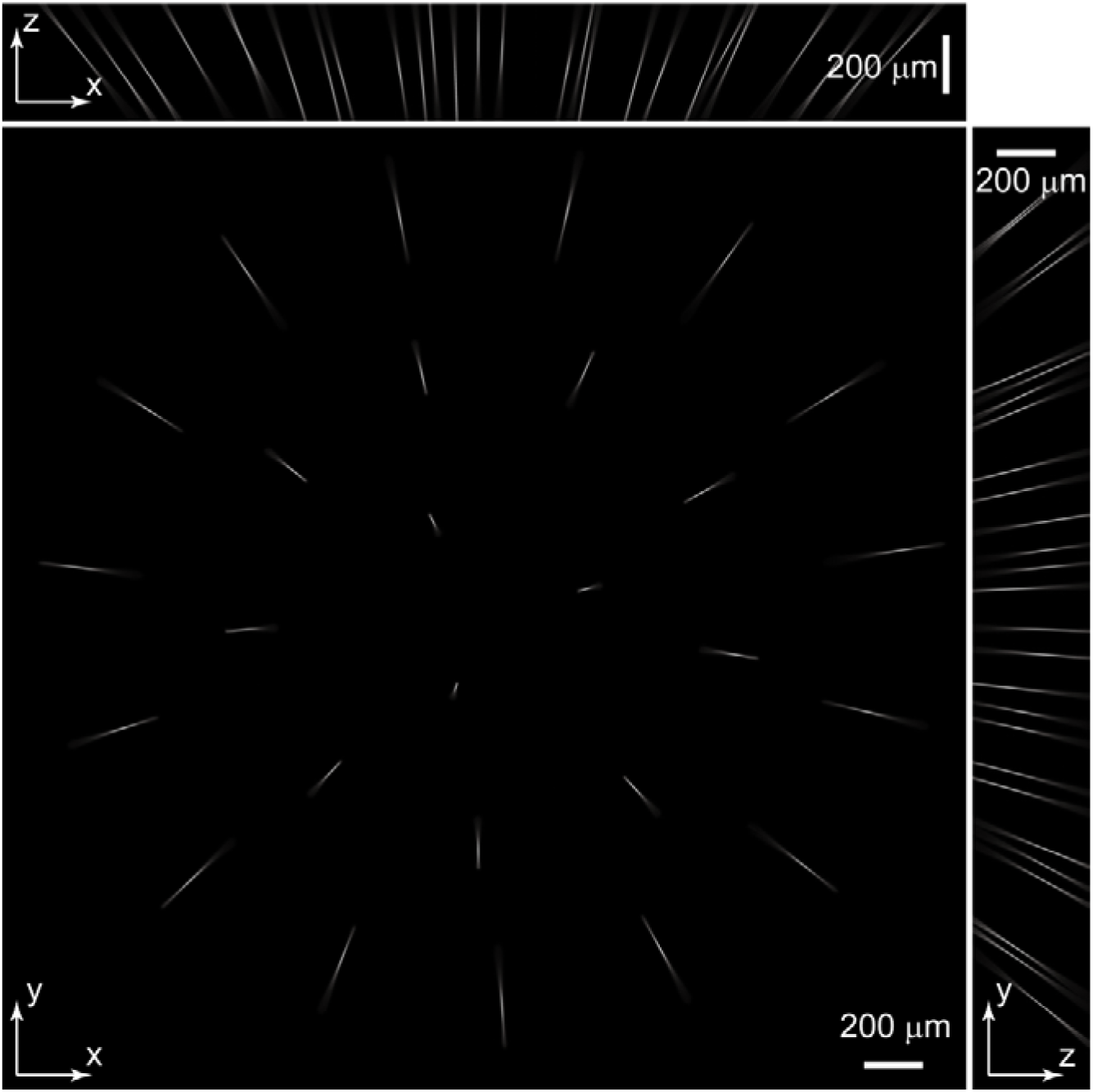
Experimentally measured PSF of the whole imaging system. Maximum intensity projections (MIPs) of the measured raw PSF stack. The stack was 2048 pixels × 2048 pixels × 200 pixels with a voxel size of 1.6 μm × 1.6 μm × 2 μm.

**Figure 1-figure supplement 3.**
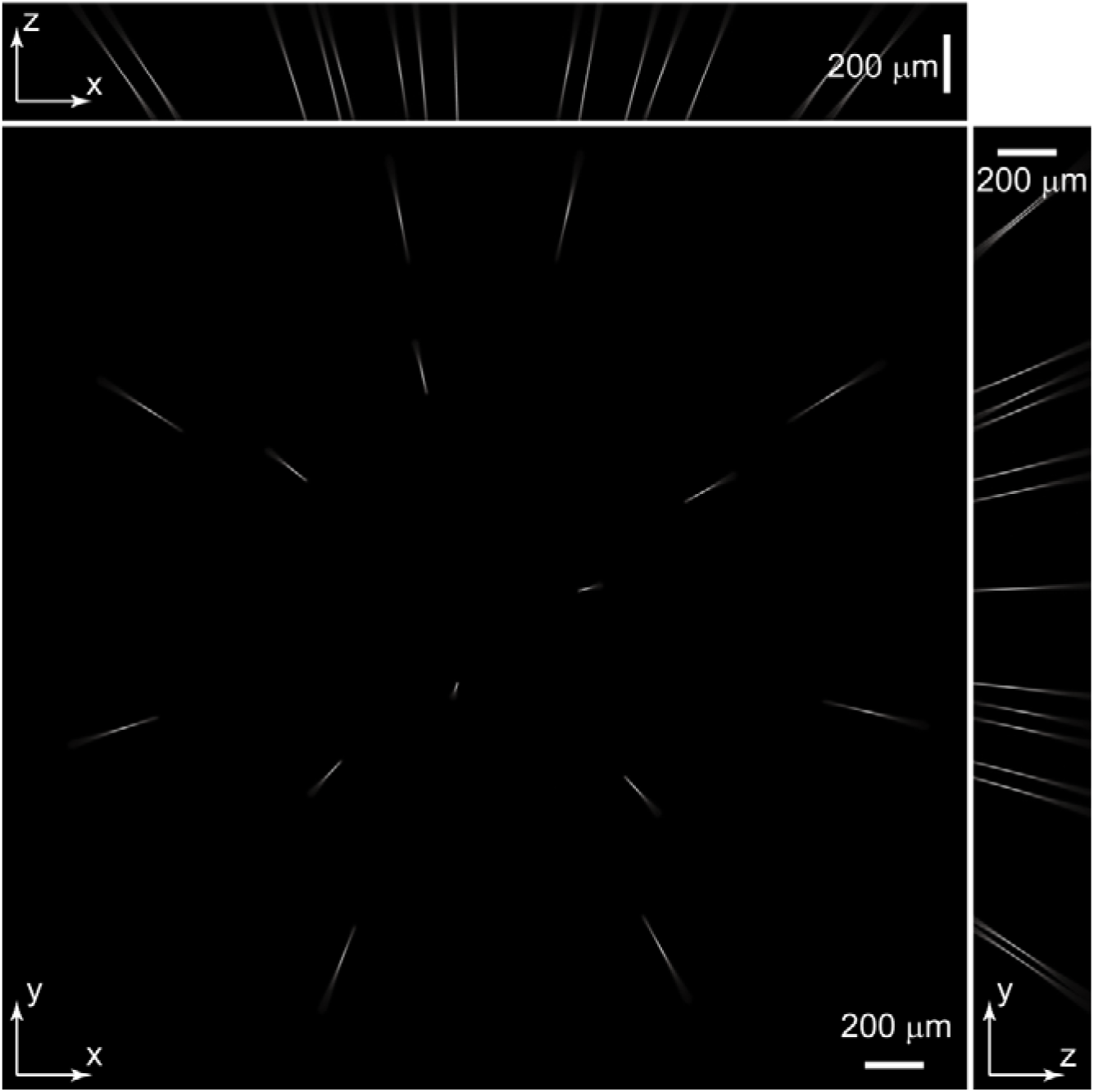
PSF of Group A micro-lenses: PSF_A. Maximum intensity projections (MIP) of PSF_A. PSF_A was extracted from experimentally measured PSF (Figure 1-figure supplement 2) according to individual micro-lens positions in group A.

**Figure 1-figure supplement 4.**
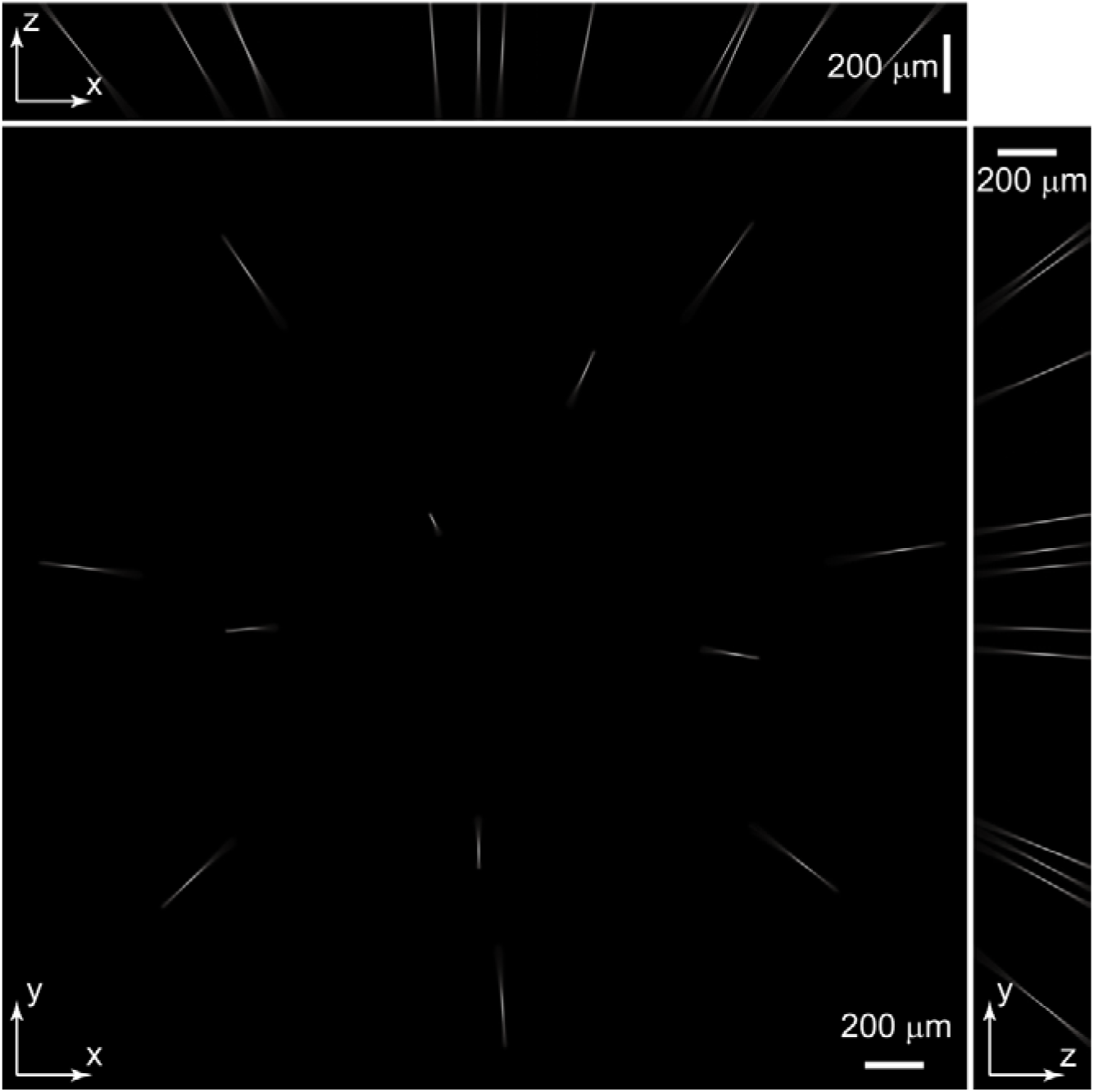
PSF of Group B micro-lenses: PSF_B. Maximum intensity projections (MIP) of PSF_B. PSF_B was extracted from experimentally measured PSFs (Figure 1-figure supplement 2) according to individual micro-lens positions in group B.

**Figure 1-figure supplement 5.**
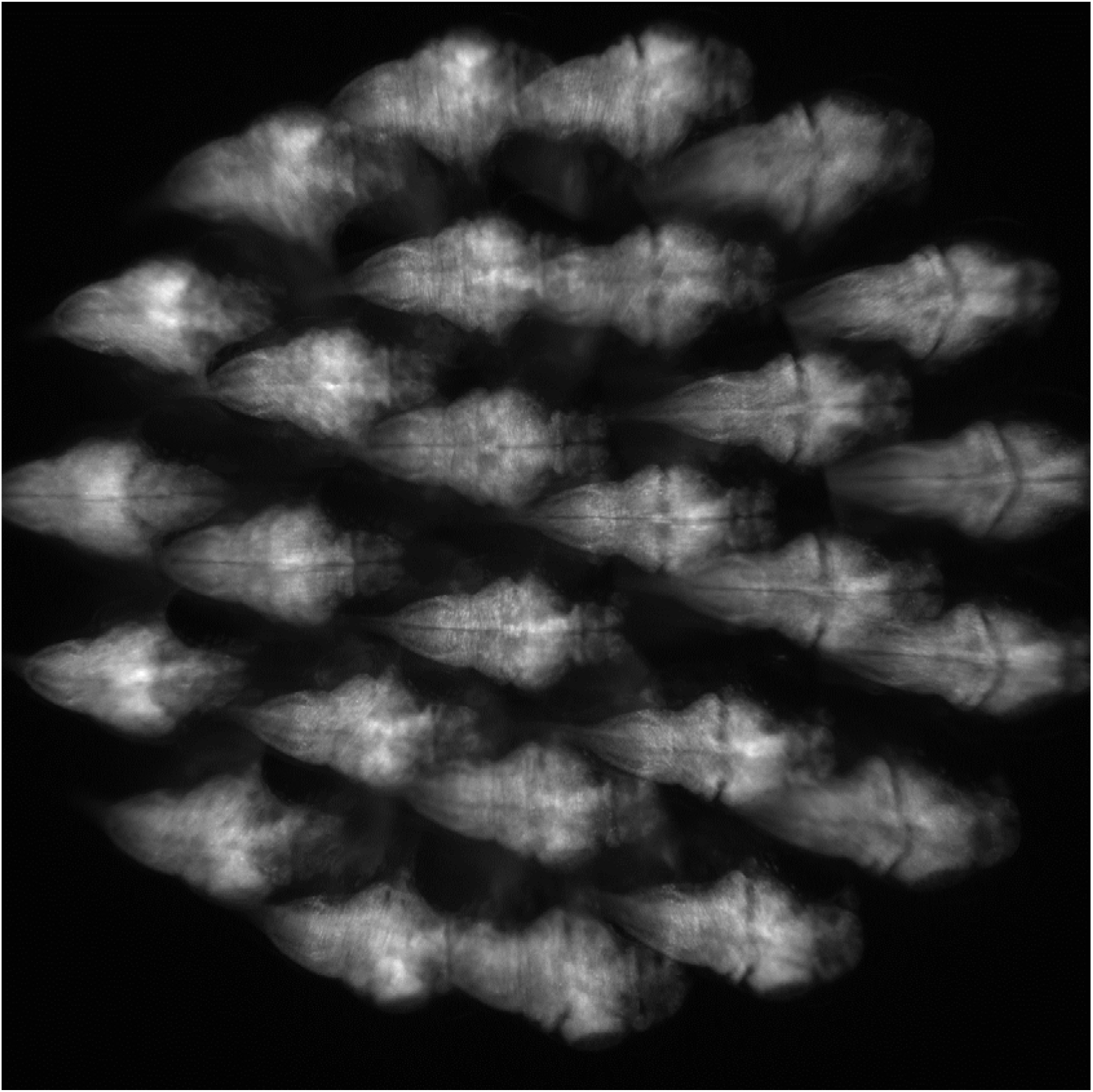
Example of camera captured raw imaging data of larval zebrafish. Raw fluorescence imaging data consisted of 27 sub-images of a larval zebrafish formed by 27 micro-lenses. Under the condition that the PSF is spatially invariant, which is satisfied apart from small aberrations, the algorithm can handle overlapping fish images.

**Figure 1-figure supplement 6.**
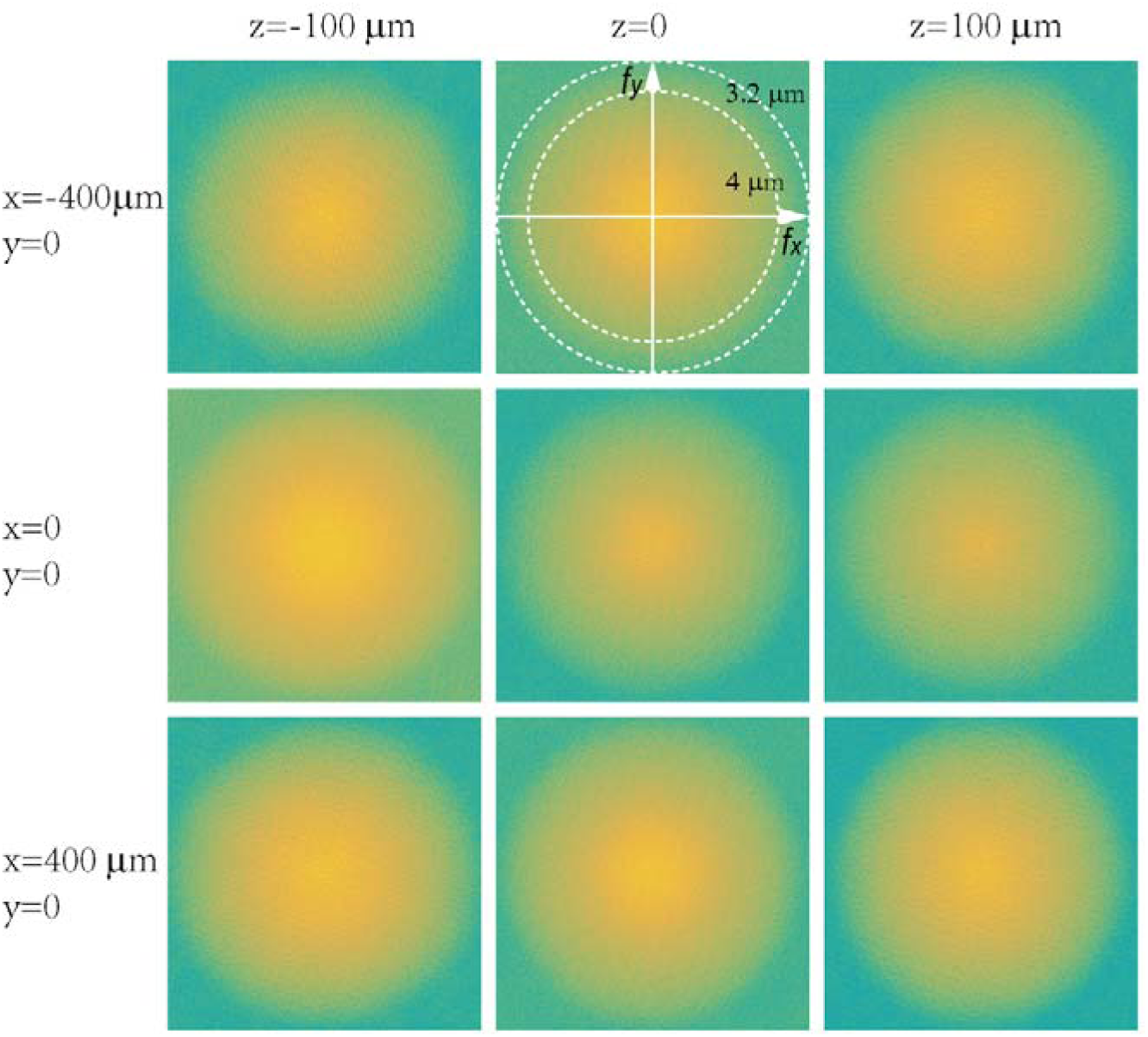
Characterization of in-plane resolution of micro-lenses. Fourier transforms of raw images of a 0.5-μm diameter fluorescent particle placed at different locations (x = −400, 0, 400 μm; z = −100, 0, 100 μm) were plotted in log scales. Dashed circles represent in-plane spatial frequency coordinates corresponding to spatial resolutions of 3.2 μm and 4 μm, respectively.

**Figure 1-figure supplement 7.**
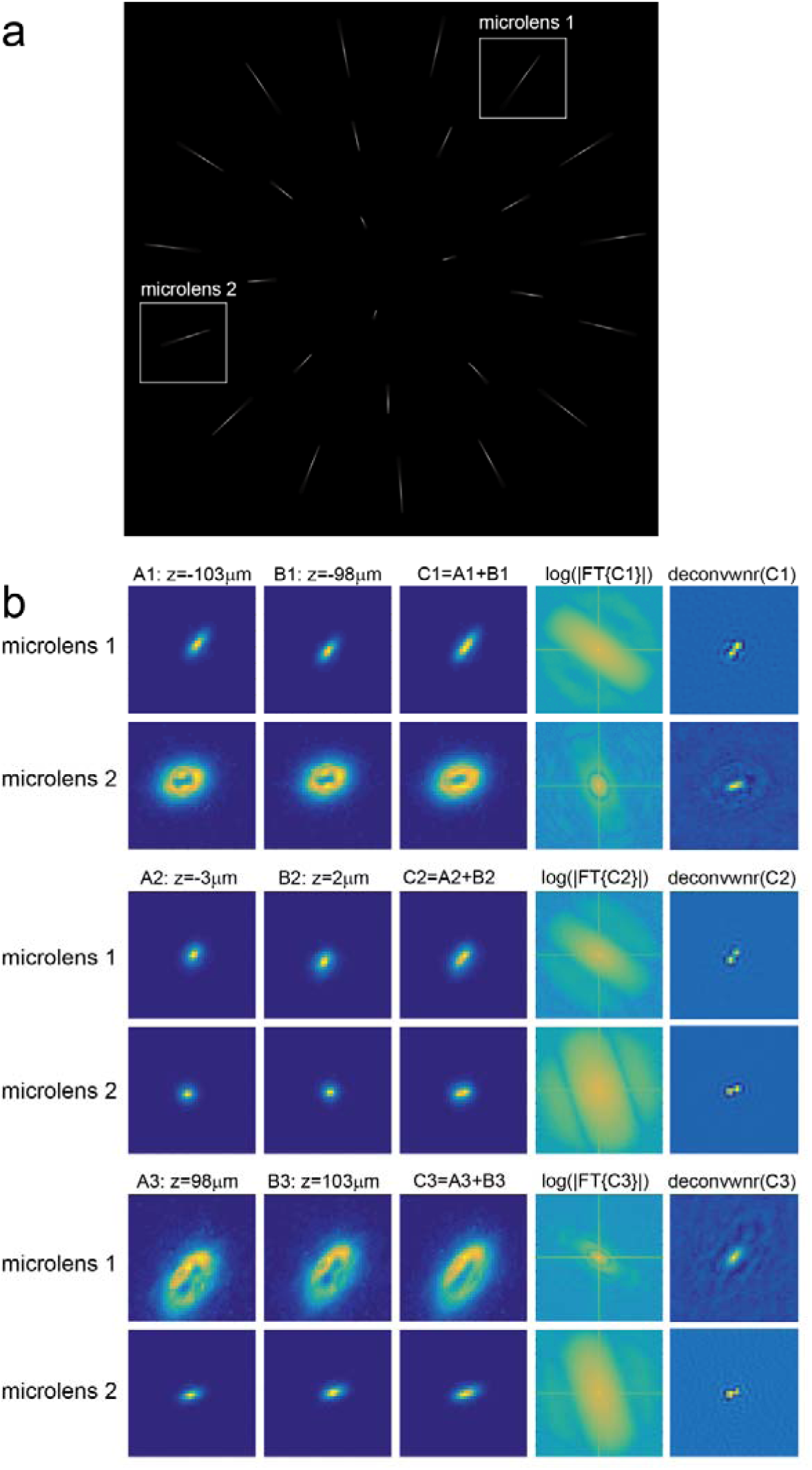
Characterization of axial resolution of XLFM afforded by individual micro-lenses. Characterization of axial resolution using a 0.5-μm diameter bright fluorescent particle. (a) Maximum intensity projection of an image stack consisting of the particle’s fluorescent images captured at different z positions. (b) Analysis of the images formed by micro-lenses 1 and 2, indicated by sub-regions in (a). The first and second columns are the particle’s fluorescent images captured at different z positions separated by 5 μm. The third column is the sum of columns 1 and 2. The fourth column is the Fourier analysis of column 3 using function: f(x) = log(|ℱ (x)|), where ℱ (x) represents the Fourier transform. The fifth column is the deconvolution of column 3 using Wiener filtering method. Experimentally measured images of the bead at different z positions (z = −100 μm, z = 0 μm and z = 100μm) are employed as PSFs to deconvolve different images (C1, C2 and C3), respectively.

**Figure 1-figure supplement 8.**
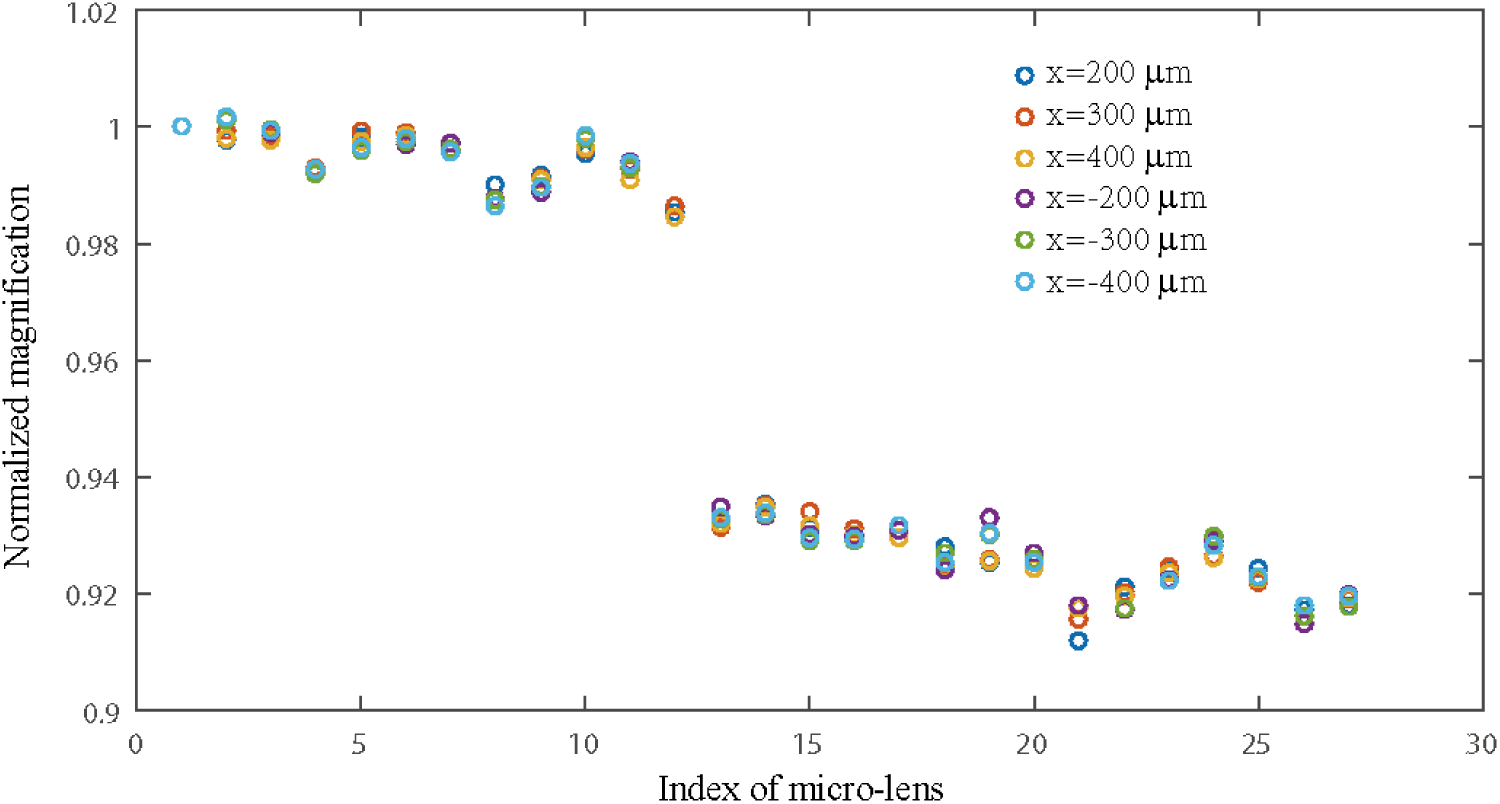
Characterization of magnification variation of micro-lenses in XLFM. Magnifications of 27 micro-lenses were measured at different locations across the field of view. A fluorescent bead originally placed at the center of the field of view (x, y, z=0) was moved to six different locations (x = 200 μm, 300 μm, 400 μm, −200 μm, −300 μm, - 400 μm, y = 0, z = 0). Six classes of the bead’s image shifts, represented by different colors, were measured. Each class consisted of 27 image shifts formed by 27 micro-lenses. Within each class, image shifts were normalized to the one from the first micro-lens. The first 12 micro-lenses and the rest formed two different groups of micro-lenses: group B and group A, consistent with Figure 1-figure supplements 3 & 4. The magnification variation of a single micro-lens across the field of view was small (< 0.3%), suggesting that the spatial invariance of individual micro-lens’ PSF was well preserved across the field of view of Ø = 800 μm. The variation across different micro-lenses within one group (A/B) was more evident (~ 2%), suggesting that the combined PSF from different micro-lenses was not perfectly spatially invariant.

**Figure 1-figure supplement 9.**
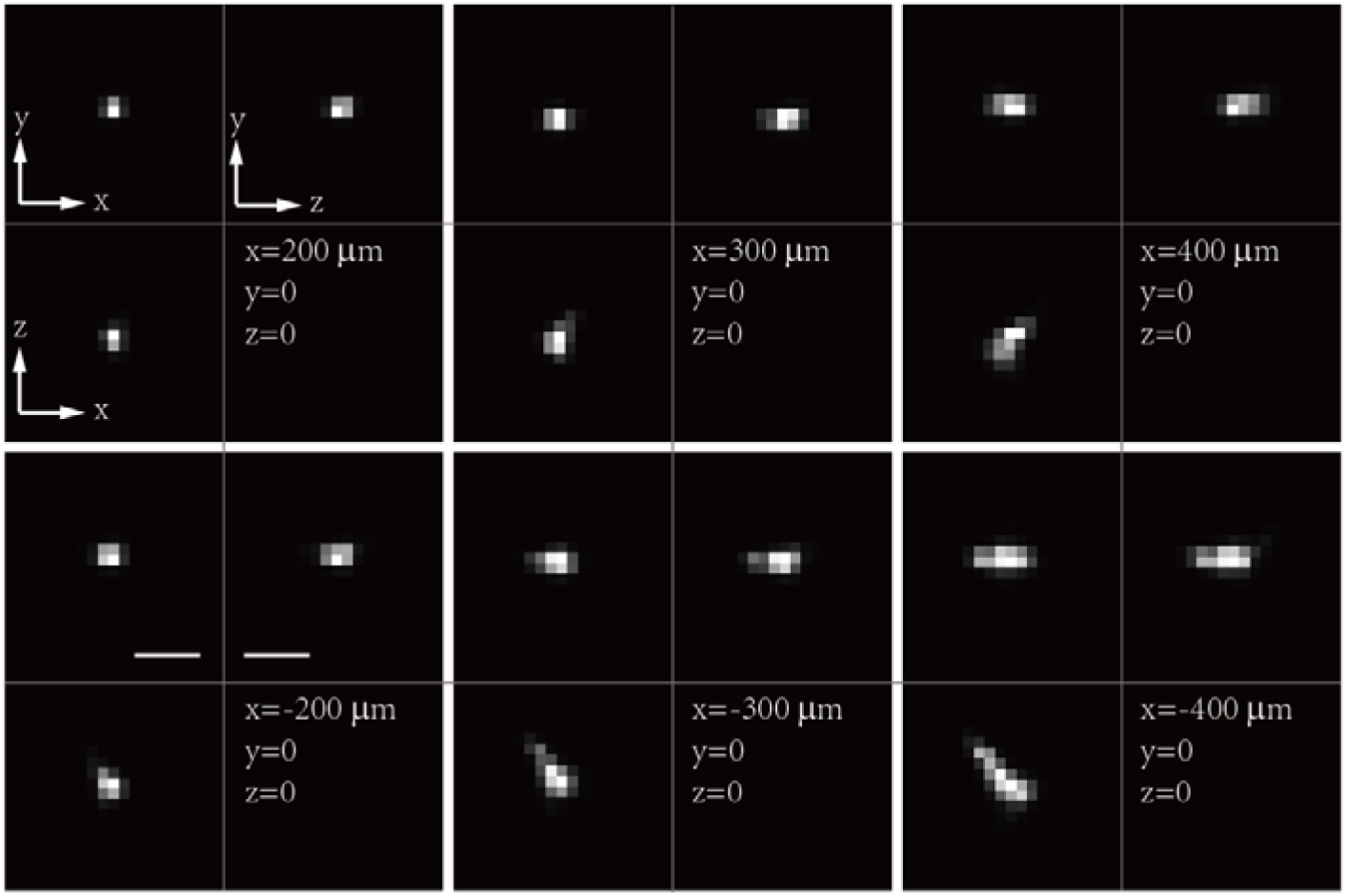
Resolution degradation due to focal length variation of micro-lenses. Maximum intensity projections (MIPs) of a reconstructed fluorescent bead positioned at different locations across the field of view. As the bead moved to the edge of the field of view, the reconstruction became distorted because the magnification variation of the micro-lenses led to spatial variance of total PSF. Scale bars are 10 μm.

**Figure 1-figure supplement 10.**
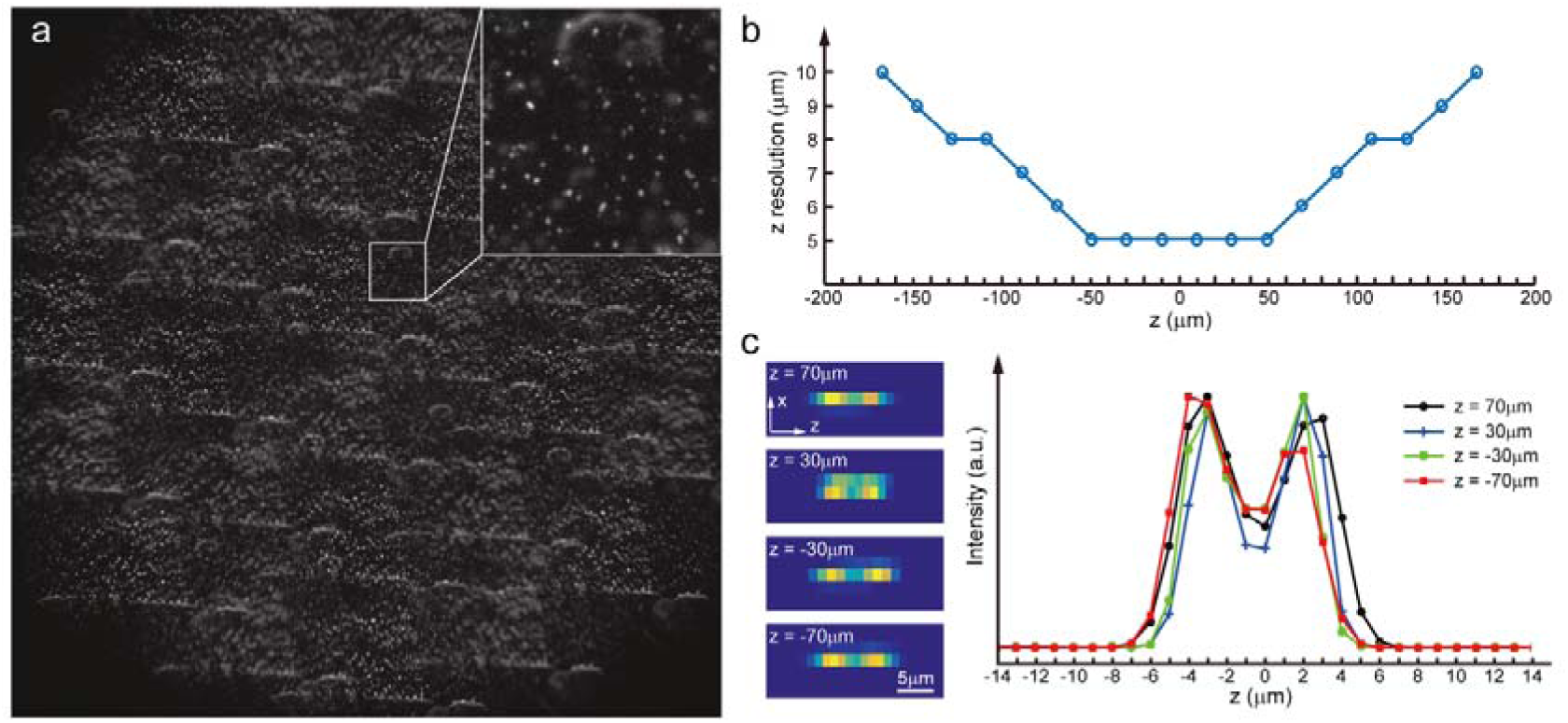
Characterization of axial resolution of XLFM at low SNR. Characterization of axial resolution using densely packed fluorescent particles (0.5 μm in diameter) at low SNR. (a) Synthetic XLFM raw image (Methods) formed by two layers of fluorescent particles with different z positions. (b) Axial resolution at different depths characterized by the minimum separation of two particles in *z*, which can be resolved using the reconstruction algorithm (Methods). (c) Left, reconstructed examples of X-Z projections of two particles located at different z positions (−70 μm, −30 μm, 30 μm, 70 μm) with different axial separations (6 μm, 5-μm, 5-μm, 6 μm); right, extracted intensity profiles of these examples.

**Figure 1-figure supplement 11.**
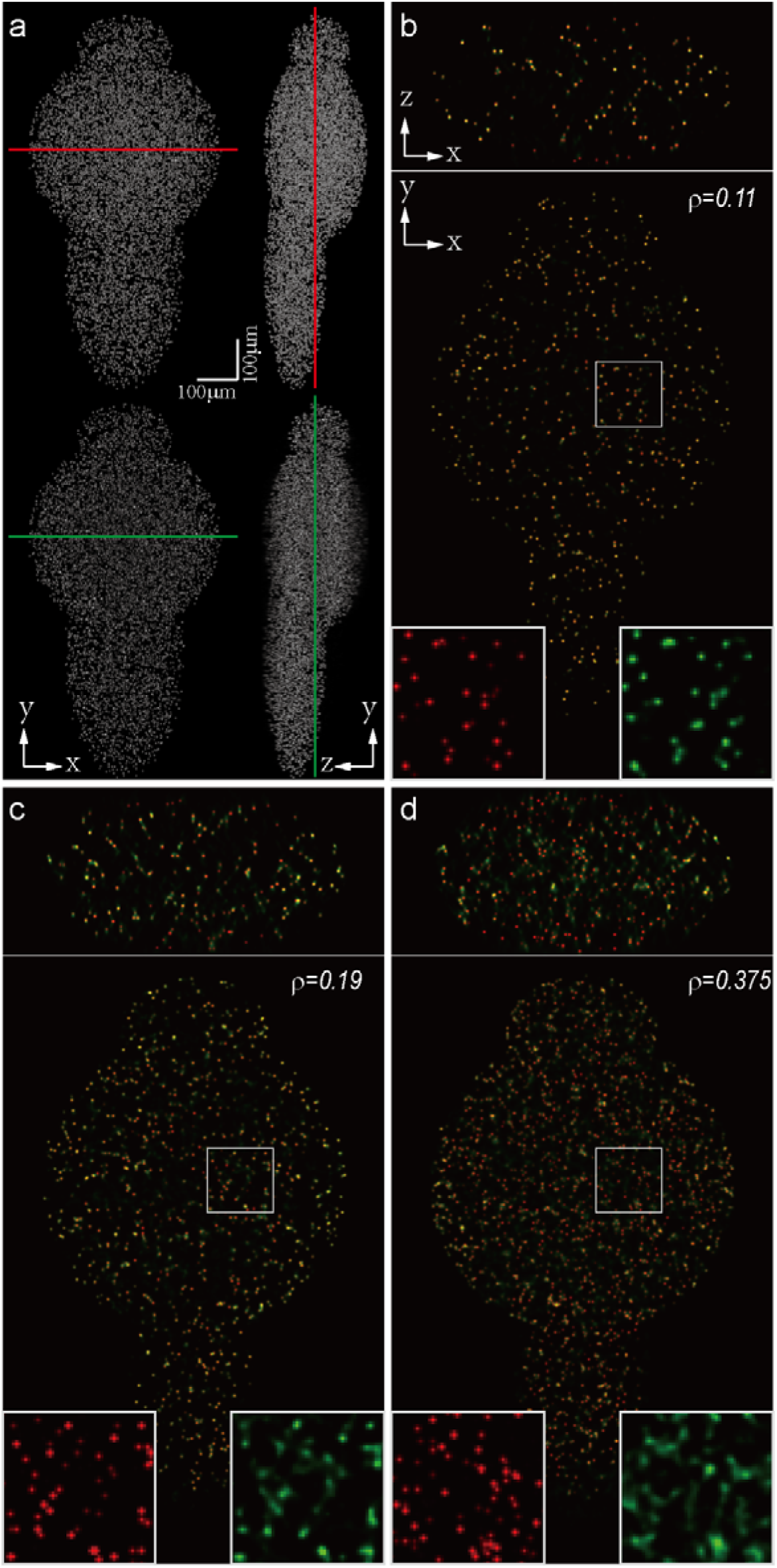
Dependence of imaging resolution on the sparseness of the sample. Characterization of the dependence of imaging resolution on the sparseness of the sample using computer simulation. (a) Maximum intensity projections (MIPs) of a numerically simulated (top) and reconstructed (bottom) larval zebrafish with randomly distributed active neurons. Red and green lines indicate positions where simulated (red) and reconstructed (green) cross-sections are compared. We assumed that the total number of neurons in the zebrafish brain is 80,000, and gradually increased the sparseness index *ρ*, the fraction of neurons activated at a given frame. (b)–(d) Characterization of the reconstruction results for different *ρ*. Insets are magnified views of rectangular regions. Red and green dots are simulated and reconstructed neurons, respectively.

**Figure 1-figure supplement 12.**
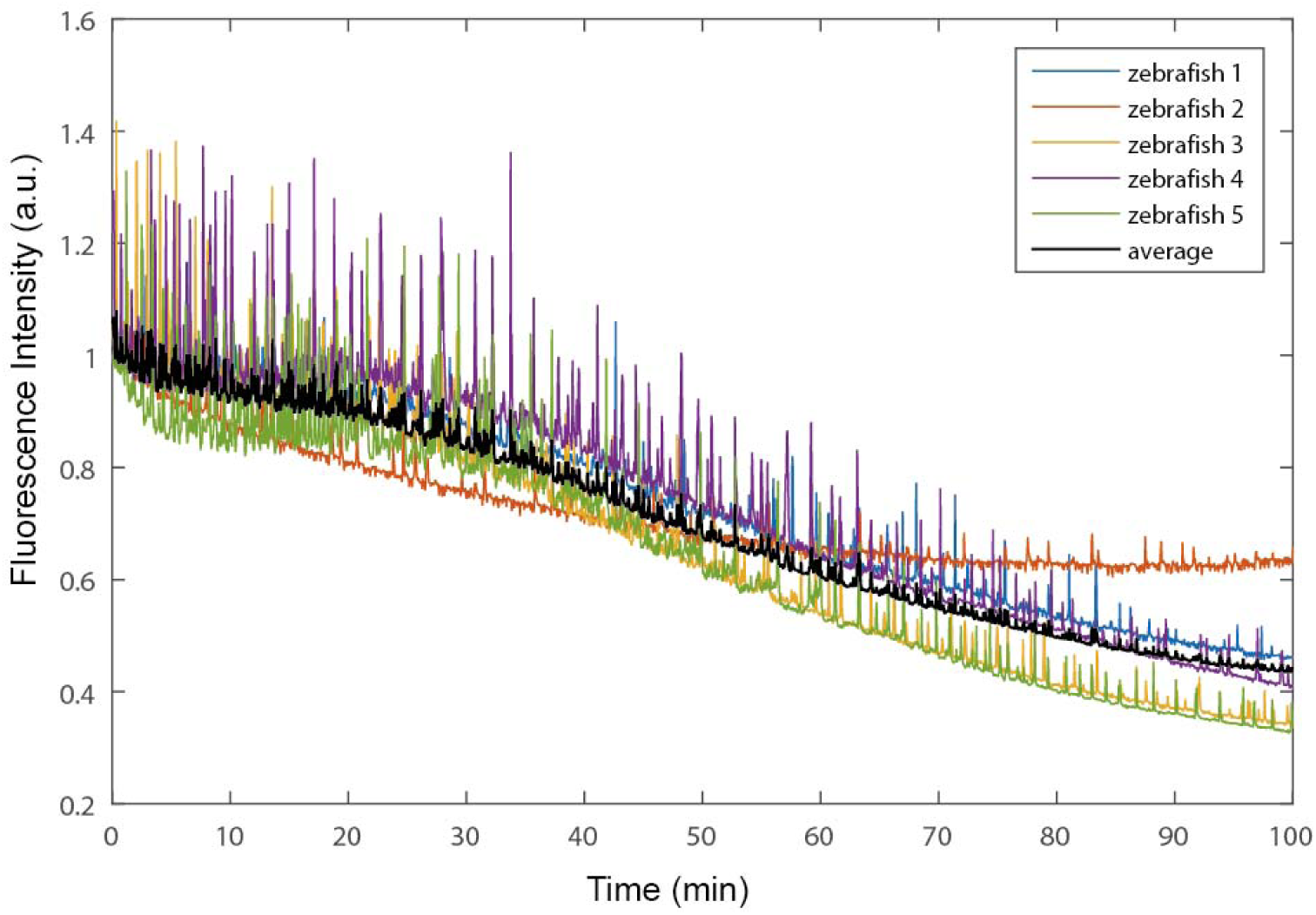
Characterization of photobleaching in fluorescence imaging by XLFM. Photobleaching was characterized by a total fluorescence intensity change of five 5 dpf zebrafish larval with nucleus-localized GCamp6f (huc:h2b-gcamp6f). Each fish was embedded in 1% agarose and continuously exposed to 2.5 mW/mm^2^ fluorescence excitation laser (488 nm) illumination. After ~100 min, corresponding to 300,000 volumes with a volume rate of 50 volumes/s, total fluorescence intensity dropped to half of that at the starting point. Random spikes corresponded to spontaneous neural activity. Fish were alive and swam normally when they were relieved from the agarose after imaging.

**Figure 2-figure supplement 1.**
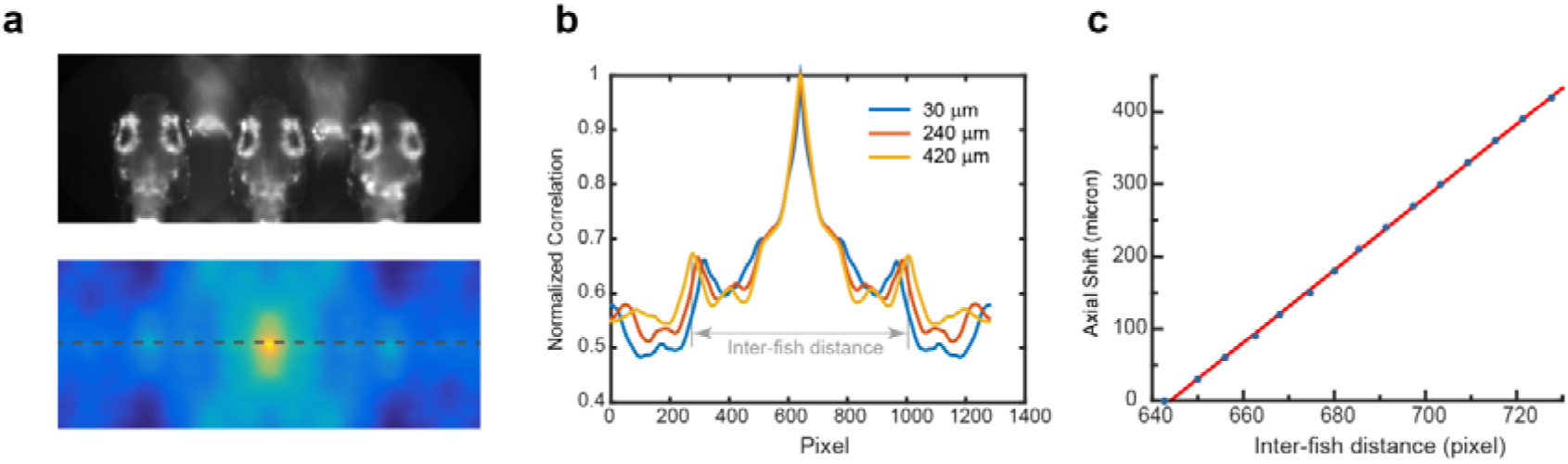
Characterization of the autofocus system. (a) Autofocus camera behind a one-dimensional lenslet array captured triplet images of the fish head (up). Its autocorrelation function was computed (bottom). (b) Central line profile of the autocorrelation function was extracted and inter-fish distance was computed as local maximums in the autocorrelation function. (c) Axial shift of the fish head, calibrated by moving the piezo at a constant interval, changed linearly (red line) with inter-fish distance.

**Source Code File 1| Computer-Aided Design files of mounting plates for micro-lenses array**

**Source Code File 2| Source code for XLFM reconstruction**

**Source Code File 3| Source code for Real-Time behavioral analysis**

## References

1 Kerr, J.N.D. and W. Denk, Imaging in vivo: watching the brain in action. Nature Reviews Neuroscience, 2008. 9(3): p. 195–205.

2 Dombeck, D.A., et al., Imaging large-scale neural activity with cellular resolution in awake, mobile mice. Neuron, 2007. 56(1): p. 43–57.

3 Wyart, C., et al., Optogenetic dissection of a behavioural module in the vertebrate spinal cord. Nature, 2009. 461(7262): p. 407–U105.

4 Boyden, E.S., et al., Millisecond-timescale, genetically targeted optical control of neural activity. Nature Neuroscience, 2005. 8(9): p. 1263–1268.

5 Zhang, F., et al., Multimodal fast optical interrogation of neural circuitry. Nature, 2007. 446(7136): p. 633–U4.

6 Chen, T.W., et al., Ultrasensitive fluorescent proteins for imaging neuronal activity. Nature, 2013. 499(7458): p. 295-+.

7 Tian, L., et al., Imaging neural activity in worms, flies and mice with improved GCaMP calcium indicators. Nature Methods, 2009. 6(12): p. 875–U113.

8 Luo, L., E.M. Callaway, and K. Svoboda, Genetic dissection of neural circuits. Neuron, 2008. 57(5): p. 634–660.

9 Friedrich, R.W., G.A. Jacobson, and P. Zhu, Circuit neuroscience in zebrafish. Curr Biol, 2010. 20(8): p. R371–81.

10 Ahrens, M.B. and F. Engert, Large-scale imaging in small brains. Curr Opin Neurobiol, 2015. 32: p. 78–86.

11 Ahrens, M.B., et al., Brain-wide neuronal dynamics during motor adaptation in zebrafish. Nature, 2012. 485(7399): p. 471–7.

12 Ahrens, M.B., et al., Whole-brain functional imaging at cellular resolution using light-sheet microscopy. Nat Methods, 2013. 10(5): p. 413–20.

13 Engert, F., The big data problem: turning maps into knowledge. Neuron, 2014. 83 (6): p. 1246–8.

14 Engert, F., Fish in the matrix: motor learning in a virtual world. Front Neural Circuits, 2012. 6: p. 125.

15 Bianco, I.H., et al., The tangential nucleus controls a gravito-inertial vestibulo-ocular reflex. Curr Biol, 2012. 22(14): p. 1285–95.

16 Bianco, I.H., A.R. Kampff, and F. Engert, Prey capture behavior evoked by simple visual stimuli in larval zebrafish. Front Syst Neurosci, 2011. 5: p. 101.

17 Patterson, B.W., et al., Visually guided gradation of prey capture movements in larval zebrafish. J Exp Biol, 2013. 216(Pt 16): p. 3071–83.

18 Trivedi, C.A. and J.H. Bollmann, Visually driven chaining of elementary swim patterns into a goal-directed motor sequence: a virtual reality study of zebrafish prey capture. Front Neural Circuits, 2013. 7: p. 86.

19 Naumann, E.A., et al., Monitoring neural activity with bioluminescence during natural behavior. Nat Neurosci, 2010. 13(4): p. 513–20.

20 Muto, A., et al., Real-time visualization of neuronal activity during perception. Curr Biol, 2013. 23(4): p. 307–11.

21 Broxton, M., et al., Wave optics theory and 3-D deconvolution for the light field microscope. Opt Express, 2013. 21(21): p. 25418–39.

22 Prevedel, R., et al., Simultaneous whole-animal 3D imaging of neuronal activity using light-field microscopy. Nat Methods, 2014. 11(7): p. 727–30.

23 Pégard, N.C., et al., Compressive light-field microscopy for 3D neural activity recording. Optica, 2016. 3(5): p. 517–524.

24 Nobauer, T., et al., Video rate volumetric Ca2+ imaging across cortex using seeded iterative demixing (SID) microscopy. Nat Meth, 2017. 14(8): p. 811–818.

25 Severi, K.E., et al., Neural control and modulation of swimming speed in the larval zebrafish. Neuron, 2014. 83(3): p. 692–707.

26 Adelson, E.H. and J.Y.A. Wang, Single Lens Stereo with a Plenoptic Camera. Ieee Transactions on Pattern Analysis and Machine Intelligence, 1992. 14(2): p. 99–106.

27 Abrahamsson, S., et al., Fast multicolor 3D imaging using aberration-corrected multifocus microscopy. Nat Meth, 2013. 10(1): p. 60–63.

28 Perwass, C. and L. Wietzke. Single lens 3D-camera with extended depth-of-field. 2012.

29 Hill, A., et al., Neurodevelopmental defects in zebrafish (Danio rerio) at environmentally relevant dioxin (TCDD) concentrations. Toxicological Sciences, 2003. 76(2): p. 392–399.

30 Ji, N., Adaptive optical fluorescence microscopy. Nat Meth, 2017. 14(4): p. 374–380.

31 Semmelhack, J.L., et al., A dedicated visual pathway for prey detection in larval zebrafish. Elife, 2014. 3.

32 Bianco, I.H. and F. Engert, Visuomotor transformations underlying hunting behavior in zebrafish. Curr Biol, 2015. 25(7): p. 831–46.

33 Venkatachalam, V., et al., Pan-neuronal imaging in roaming Caenorhabditis elegans. Proc Natl Acad Sci U S A, 2016. 113(8): p. E1082–8.

34 Nguyen, J.P., et al., Whole-brain calcium imaging with cellular resolution in freely behaving Caenorhabditis elegans. Proc Natl Acad Sci U S A, 2016. 113(8): p. E1074–81.

35 Berg, H.C., How to Track Bacteria. Review of Scientific Instruments, 1971. 42 (6): p. 868-&.

36 Ng, R., et al., Light Field Photography with a Hand-held Plenoptic Camera. Stanford Tech Report 2005.

37 Levoy, M., et al., Light field microscopy. ACM Trans. on Graphics (Proc. SIGGRAPH), 2006. 25: p. 924–934.

38 Niell, C.M. and M.P. Stryker, Modulation of visual responses by behavioral state in mouse visual cortex. Neuron, 2010. 65(4): p. 472–9.

39 Maimon, G., A.D. Straw, and M.H. Dickinson, Active flight increases the gain of visual motion processing in Drosophila. Nat Neurosci, 2010. 13(3): p. 393–9.

40 Chiappe, M.E., et al., Walking modulates speed sensitivity in Drosophila motion vision. Curr Biol, 2010. 20(16): p. 1470–5.

41 Pearson, K.G., Proprioceptive regulation of locomotion. Current Opinion in Neurobiology, 1995. 5(6): p. 786–791.

42 Bell, C.C., An Efference Copy Which Is Modified by Reafferent Input. Science, 1981. 214(4519): p. 450–453.

43 Portugues, R. and F. Engert, Adaptive locomotor behavior in larval zebrafish. Front Syst Neurosci, 2011. 5: p. 72.

44 Portugues, R., et al., Whole-brain activity maps reveal stereotyped, distributed networks for visuomotor behavior. Neuron, 2014. 81(6): p. 1328–43.

45 Hromadka, T., M.R. Deweese, and A.M. Zador, Sparse representation of sounds in the unanesthetized auditory cortex. PLoS Biol, 2008. 6(1): p. e16.

46 Buzsaki, G. and K. Mizuseki, The log-dynamic brain: how skewed distributions affect network operations. Nat Rev Neurosci, 2014. 15(4): p. 264–78.

47 Olshausen, B.A. and D.J. Field, Emergence of simple-cell receptive field properties by learning a sparse code for natural images. Nature, 1996. 381(6583): p. 607–9.

48 Olshausen, B.A. and D.J. Field, Sparse coding of sensory inputs. Curr Opin Neurobiol, 2004. 14(4): p. 481–7.

49 Coombs, S., et al., The lateral line system. Springer handbook of auditory research,. 2014, New York: Springer. xiv, 347 pages.

50 Liao, J.C., Organization and physiology of posterior lateral line afferent neurons in larval zebrafish. Biol Lett, 2010. 6(3): p. 402–5.

51 St-Pierre, F., et al., High-fidelity optical reporting of neuronal electrical activity with an ultrafast fluorescent voltage sensor. Nat Neurosci, 2014. 17(6): p. 884–9.

52 Ji, N., D.E. Milkie, and E. Betzig, Adaptive optics via pupil segmentation for high-resolution imaging in biological tissues. Nat Meth, 2010. 7(2): p. 141–147.

53 Wang, K., et al., Rapid adaptive optical recovery of optimal resolution over large volumes. Nat Meth, 2014. 11(6): p. 625–628.

54 Wang, K., et al., Direct wavefront sensing for high-resolution in vivo imaging in scattering tissue. Nature Communications, 2015. 6: p. 7276.

